# Sufficient levels of BECLIN1 are required for intestinal epithelial cell homeostasis and protection against unwanted intestinal inflammation

**DOI:** 10.1101/2025.04.29.651340

**Authors:** Juliani Juliani, Sharon Tran, Tiffany J. Harris, Sarah L. Ellis, Aysha H Al-Ani, Komal M. Patel, Sam N. Young, Marco Evangelista, David Baloyan, Camilla M. Reehorst, Rebecca Nightingale, Laura J. Jenkins, Peter De Cruz, Kinga Duszyc, Benjamin T. Kile, Alpha S. Yap, John M. Mariadason, Britt Christensen, André L. Samson, James M. Murphy, Walter D. Fairlie, Erinna F. Lee

**Author notes:** Corresponding authors: E.F. Lee,; W.D. Fairlie.

## Abstract

The prototypical autophagy regulator BECLIN1, orchestrates both autophagic and endocytic trafficking, and its homozygous deletion in the intestinal epithelium leads to intestinal disruption bearing similarities to inflammatory bowel disease (IBD). However, complete loss of BECLIN1 is rare in human disease. To model the effects of a partial reduction in BECLIN1, we examined mice with a monoallelic deletion of *Becn1* in the intestinal epithelium, which decreased BECLIN1 protein levels by approximately 50%. Unlike the fatal phenotype following homozygous *Becn1* deletion, the heterozygous mice were grossly normal, though they presented with significantly shorter small intestines. Gastrointestinal organoids derived from the mice also displayed disrupted endocytic trafficking and altered cytoskeletal dynamics resulting in mislocalisation of junctional proteins such as E-CADHERIN. In the mice, these changes were associated with impaired goblet cell function and maturation accompanied by abnormal mucin production, composition and secretion, resulting in compromised mucus barrier integrity. The mice also exhibited heightened susceptibility to dextran sulfate sodium (DSS)-induced colitis, emphasising the critical role of BECLIN1 in mediating protection against unregulated inflammation. Investigation of BECLIN1 levels in biopsies from patients with inflammatory bowel disease (IBD) revealed changes in its expression across distinct states of inflammation with reduced levels observed in highly inflamed regions in some patients, consistent with the gastrointestinal defects seen in mice with reduced BECLIN1. Hence, this study provides crucial insights into the multifaceted importance of BECLIN1 sufficiency in the intestinal epithelium to prevent pathophysiology. This highlights the potential of BECLIN1 as a biomarker and therapeutic target for IBD management.

## INTRODUCTION

Genome-wide association studies have identified polymorphisms in key autophagy regulators, such as *ATG16L1* and *IRGM*, as strongly associated with the development of inflammatory bowel disease (IBD) ^1–11^. These findings highlight the critical role of autophagy in maintaining intestinal homeostasis and suggest that disruptions to autophagy-related processes contribute to IBD pathogenesis. Functional studies using both mouse and cell-based models with targeted manipulations of autophagy regulators have uncovered multiple mechanisms and cellular compartments whereby autophagy exerts a protective role. These include intrinsic mechanisms within intestinal epithelial cells (IECs), such as maintaining barrier integrity and cell survival ^12–23^, as well as extrinsic mechanisms mediated by immune cells, including regulation of inflammatory responses and microbial homeostasis ^24–29^. This body of evidence highlights the many functions of autophagy in intestinal homeostasis and underscores the multifactorial nature of IBD pathogenesis, where disruptions in autophagy pathways across different cell types converge to drive disease development and progression.

Functional studies have recently highlighted the critical role of the prototypical autophagy regulator BECLIN1 in intestinal homeostasis. Constitutive activation of autophagy *via* BECLIN1 alleviates endoplasmic reticulum (ER) stress in goblet cells to produce a thicker and less penetrable mucus barrier ^18^. This, in turn, alters the gut microbiota to protect against both chemical- and infection-driven inflammation. We, and others, have also implicated an equally critical role for the alternate function of BECLIN1 in endocytic trafficking in intestinal homeostasis. This is exerted through the maintenance of barrier integrity by regulating the localisation of junctional proteins, cytoskeletal reorganisation and epithelial remodelling ^18, 30, 31^. Indeed, we recently reported that complete loss of BECLIN1 in the intestinal epithelium is rapidly fatal due to a severe enterocolitis phenotype ^31^ which was associated with morphological disruption along the length of the small intestine with excessive epithelial cell death, disruption to specialised IEC functions, increased inflammation and a breakdown in barrier integrity.

In diseases, such as cancer and neurodegenerative diseases, changes in BECLIN1 expression levels are observed rather than being caused by mutations ^32–39^. Such changes can be driven by various factors including interactions with other proteins, proteolytic cleavage, epigenetic regulation, or due to the monoallelic deletion of the *Becn1* gene ^40–46^. Despite strong functional data supporting a multi-faceted role for BECLIN1 in the maintenance of intestinal homeostasis, to date, BECLIN1 has not been identified through IBD GWAS studies as harbouring disease-associated single nucleotide polymorphisms (SNPs). However, there is emerging evidence implicating indirect IEC-intrinsic and -extrinsic mechanisms that regulate BECLIN1 levels, whereby decreased BECLIN1 levels lead to dysfunctional intestinal homeostasis and exacerbation of colitis or IBD pathogenesis. For example, the cytoplasmic pattern recognition receptor, cyclic GMP-AMP synthase (cGAS) that senses double-stranded DNA, directly interacts with BECLIN1 to promote autophagy in IECs and prevent cell death ^47^. Consequently, cGAS deficiency and, in turn, reduced BECLIN1 levels lead to worsened colitis. Similarly, the m^6^A nuclear reader, YTHDC1, which inhibits the macrophage-mediated inflammatory response by stabilising *Becn1* mRNA, and enhancing autophagy, is significantly downregulated in DSS-induced mouse models of colitis and in faecal microbiota from IBD patients ^46, 48^.

Here, we explore the mechanisms by which reduced BECLIN1 levels compromise gut mucosal immunity and increase susceptibility to inflammation, by impacting goblet cells. By dissecting the roles of BECLIN1 in regulating endocytic trafficking, cytoskeletal organisation and mucin secretion, this study provides new insights into how BECLIN1 contributes to epithelial dysfunction and potentially IBD pathogenesis.

## RESULTS

### Generation of mice where monoallelic deletion of *Becn1* can be induced

The majority of disease-related molecular changes associated with BECLIN1, including loss of heterozygosity and epigenetic or post-translational modifications, lead to a reduction in its levels. We previously showed that homozygous *Becn1* deletion, specifically in the intestinal epithelium, is fatal in adult mice within 7 days, due to a severe enterocolitis phenotype bearing features similar to IBD ^31^. To circumvent this lethality, and more closely reflect a disease setting where BECLIN1 levels are reduced rather than eliminated, we generated a model in which monoallelic deletion of *Becn1* can be induced specifically in the intestinal epithelium of adult mice. This was achieved using mice heterozygous for the *Becn1* floxed allele (*Becn1^fl/+^)* carrying the *Vil1-Cre^ERT^*^2^ transgene (*Becn1^fl/+^*;*Vil1-CreERT2^Cre/+^*), which drives Cre recombinase expression specifically in the intestinal epithelium. Littermates that did not harbour the floxed allele (*Becn1^+/+^*) but were positive for *Vil1-CreERT2^Cre/+^* (*Becn1^+/+^*;*Vil1-CreERT2^Cre/+^*) were used as controls throughout this study. Intraperitoneal Tamoxifen injections successfully induced monoallelic *Becn1* deletion in the small intestine (duodenum, jejunum and ileum) and colon of *Becn1^ff/+^;Vil1-CreERT2^Cre/+^* mice genomically and at a protein level, where there was an approximately 50% reduction in BECLIN1 protein levels (Figure 1A, Supplementary Figure 1A). The resulting Tamoxifen-treated *Becn1^+/+^*;*Vil1-CreERT2^Cre/+^* and *Becn1^fl/+^*;*Vil1-CreERT2^Cre/+^* mice will hereafter be referred to as *Becn1^+/+^* and *Becn1^+/-^* mice respectively.

**Figure 1.**
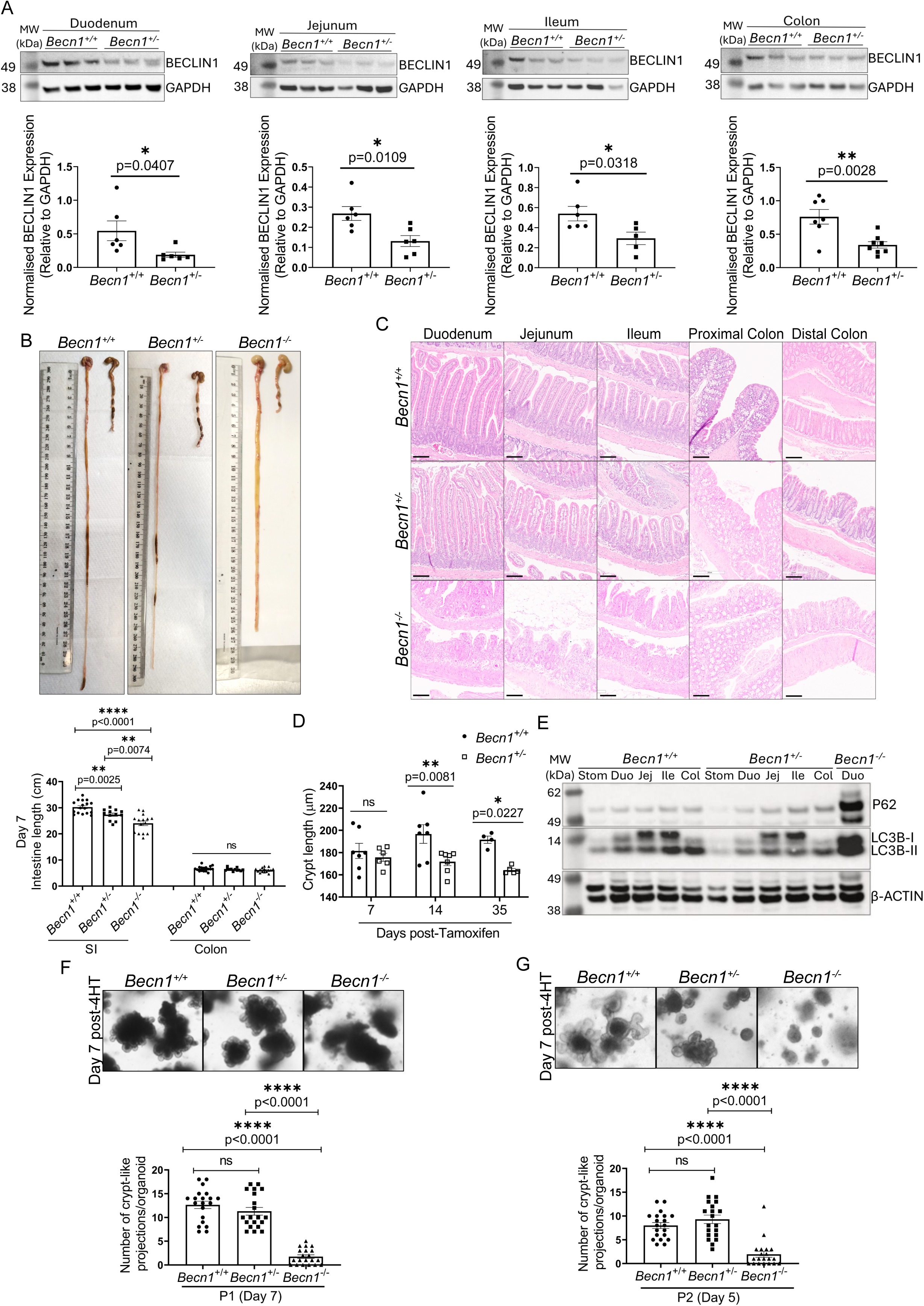
Monoallelic *Becn1* deletion induces mild intestinal epithelial alterations under basal conditions that is distinct from the severe fatal enteritis defects observed with homozygous *Becn1* deletion. (A) Representative Western blots showing BECLIN1 levels in the duodenum, jejunum, ileum and colon from a subset of the total mice analysed, along with their respective densitometry quantification below each immunoblot. GAPDH was used as a loading control for all Western blots. Data represent a minimum of *n =* 5 animals per genotype from three independent experiments. **(B)** Representative images of intestinal tracts of *Becn1^+/+^*, *Becn1^+/-^* and *Becn1^-/-^*mice along with the measurements of intestinal length at Day 7 post Tamoxifen administration. Data represent *n* > 12 animals per genotype from *n* > 3 independent experiments **(C)** HCE-stained FFPE sections of the intestinal tracts from *Becn1^+/+^, Becn1^+/-^* and *Becn1^-/-^* mice. Scale bars represent 100 μm. Data represent *n* > 6 biologically independent mice of each genotype from *n* = 3 independent experiments. **(D)** Quantification of distal colon crypt length in *Becn1^+/+^* and *Becn1^+/-^* mice at days 7, 14 and 35 post-tamoxifen administration. Only visible full-length crypts were included in the analysis, with a minimum of *n >* 5 crypts measured per mouse and averaged. Data represent *n >* 4 animals per genotype from *n =* 3 independent experiments. **(E)** Western blot showing markers of autophagy flux across different sections of the gastrointestinal tract in *Becn1^+/+^* and *Becn1^+/-^* mice. The duodenum of *Becn1^-/-^* mice was included as a positive control for defective autophagy. β-ACTIN was used as a loading control. **(F)** Representative phase contrast microscopy images of *Becn1^+/+^*, *Becn1^+/-^* and *Becn1^-/-^* organoids at Day 7 post-4-HT treatment and **(G)** re-passaged surviving *Becn1^+/+^*, *Becn1^+/-^* and *Becn1^-/-^* organoids at Day 5, along with respective quantification of the number of budding crypt-like projections per organoid. Data in (F) and (G) were obtained from *n =* 20 organoids per genotype across *n =* 3 independent experiments. Graphs show the mean ± S.E.M. Statistical significance was determined by unpaired (Student’s) t-test, except in (F) and (G) where ordinary one-way ANOVA was used. SI: small intestine. Stom: stomach. Duo: duodenum. Jej: jejunum. Ile: ileum. Col: Colon. MW: molecular weight. 4-HT: 4-hydroxytamoxifen. ns: not significant (p > 0.05)

### Mice with reduced BECLIN1 levels in the intestinal epithelium present with shortened small intestines and reduced crypt lengths but no other clinical abnormalities

At 7 days post gene deletion, there were no significant differences between the body weights of *Becn1^+/+^* and *Becn1^+/-^* mice (Supplementary Figure 1B), contrasting the significant weight loss observed when *Becn1* is homozygously deleted ^31^. Despite appearing grossly normal, *Becn1^+/-^*mice exhibited significantly shorter small intestines compared to *Becn1^+/+^* controls, whilst their colon lengths remained unchanged (Figure 1B). Staining with HCE showed healthy-looking epithelium of the small and large intestines with intact morphology in both *Becn1^+/+^* and *Becn1^+/-^*animals (Figure 1C). There were no significant differences in crypt length (Figure 1D) or width (Supplementary Figure 1C) in the distal colon between *Becn1^+/+^* and *Becn1^+/-^* mice at this time point. Interestingly however, at 14- and 35-days post gene deletion, we observed a significant reduction in crypt length in *Becn1^+/-^*compared to *Becn1^+/+^*mice (Figure 1D). This suggests a reduction in the volume fraction of the colonic crypts with age when BECLIN1 levels are reduced^49^. Aging of the *Becn1^+/-^* mice for up to a month also resulted in a significantly shorter small intestinal tract (Supplementary Figure 1D) though there was no significant body weight loss, and the entire intestinal tract appeared morphologically normal (Supplementary Figure 1B, E). All these observations in the *Becn1^+/-^*mice differed significantly from the intestinal epithelium of *Becn1^-/-^* mice where the small intestine was significantly shortened and appeared swollen and lytic, with pronounced villus stunting along the entire length, within 7 days post gene deletion as we have previously reported (Figures 1B, C) ^31^. Notably, the requirement to euthanise *Becn1*^⁻/⁻^ mice within 7 days of inducing homozygous deletion may have precluded the emergence of any detectable phenotype in the colon in those mice. Overall, apart from the reduction in small intestinal length and colonic crypt length, heterozygous deletion of *Becn1* in adult mice in contrast to homozygous deletion ^31^, does not result in dramatic overt and spontaneous intestinal abnormalities under basal conditions up to one month post gene deletion.

### Intestinal epithelial cells with reduced BECLIN1 levels displayed no observable disruption to basal autophagy flux, cell differentiation, and proliferation

We next assessed the functional consequences of heterozygous *Becn1* loss in the intestinal epithelium. In the first instance, we investigated the impact on autophagy in isolated IECs. Heterozygous *Becn1* deletion resulted in minimal changes to basal autophagy flux, as assessed by total P62 and LC3B levels that were comparable to wild-type IECs (Figure 1E). This contrasted *Becn1^-/-^* IECs where BECLIN1 protein levels were reduced by greater than 50% ^31^ and an obvious impairment in autophagy flux was evident due to the pronounced accumulation of total P62 and LC3B levels (Figure 1E) ^31^.

We previously identified an essential role for BECLIN1 in ensuring an intact endocytic trafficking pathway in intestinal-derived organoids ^30, 31^. To facilitate investigation of whether monoallelic loss of BECLIN1 induces similar defects in endocytic trafficking, we isolated and expanded intestinal crypts from *Becn1^+/+^*; *Becn1^fl/+^*; and *Becn1^ff/ff^*;*Vil1-CreERT2^Cre/+^* mice to generate intestinal epithelial organoid cultures. Treatment with the Tamoxifen metabolite, 4-hydroxytamoxifen (4-HT), resulted in BECLIN1 deletion in *Becn1^ff/ff^*;*Vil1-CreERT2^Cre/+^* (*Becn1^-/-^*, 100% deletion) and *Becn1^fl/+^*;*Vil1-CreERT2^Cre/+^* (*Becn1^+/-^*, approximately 50% deletion) organoids, but not in *Becn1^+/+^*;*Vil1-CreERT2^Cre/+^* (*Becn1^+/+^*) IECs (Supplementary Figure 2A). Consistent with observations in IECs isolated from mice, monoallelic deletion of *Becn1* in intestinal organoids had minimal impact on basal autophagy flux in contrast to when *Becn1* was homozygously deleted (Supplementary Figure 2B). At 7 days post 4-HT addition, whilst *Becn1^-/-^*organoids showed significantly reduced budding crypt formation (Figure 1F) ^31^, *Becn1^+/-^*organoids did not display any observable morphological defects, with similar numbers of crypt-like projections forming per organoid as *Becn1^+/+^* organoids indicative of normal IEC proliferation and differentiation (Figure 1F). This was further confirmed *in vivo,* as intestinal tissues from *Becn1^+/-^* mice showed no significant difference in IEC proliferation, including in the crypt base, compared to *Becn1^+/+^*mice, as assessed by Ki67 staining (Supplementary Figure 1F). Re-passaging of *Becn1^+/+^* and *Becn1^+/-^* also continuously yielded mature organoids by Day 5, with similar numbers of budding crypts, emphasising minimal defects in proliferation and differentiation of *Becn1^+/-^* IECs (Figure 1G). In contrast, re-passaging of any surviving *Becn1^-/-^*organoids subsequently failed to grow viable crypt structures, evidenced by significantly fewer crypt-like projections at Day 5 compared to both *Becn1^+/+^* and *Becn1^+/-^* organoids (Figure 1G). Hence, our results show that reduced BECLIN1 levels do not significantly impact basal autophagy or the survival, differentiation and proliferation of IECs.

### *Becn1^+/-^* IECs display changes early in the endocytic trafficking pathway and the distribution of E-CADHERIN

We next investigated if heterozygous loss of BECLIN1 influences endocytic trafficking as occurs following its homozygous loss ^30, 31, 50–54^. Similar to what we have previously reported following homozygous loss of *Becn1*, there was a small but significant increase in the number of RAB5^+ve^ early endosomes in the cytoplasm of *Becn1^+/-^* IECs (Figures 2A, B, Supplementary Figure 3) ^31^. However, unlike in *Becn1*^-/-^ organoids, the RAB5^+ve^ puncta were not aberrantly enlarged (Figure 2C) ^31^. In addition, whilst there was also significantly decreased RAB5 staining along the apical and lateral membranes of *Becn1^-/-^* IECs compared to *Becn1^+/+^* cells, there were no significant changes in the amount of RAB5 localised on the plasma membrane of *Becn1^+/-^* IECs (Figures 2A, D, E).

**Figure 2.**
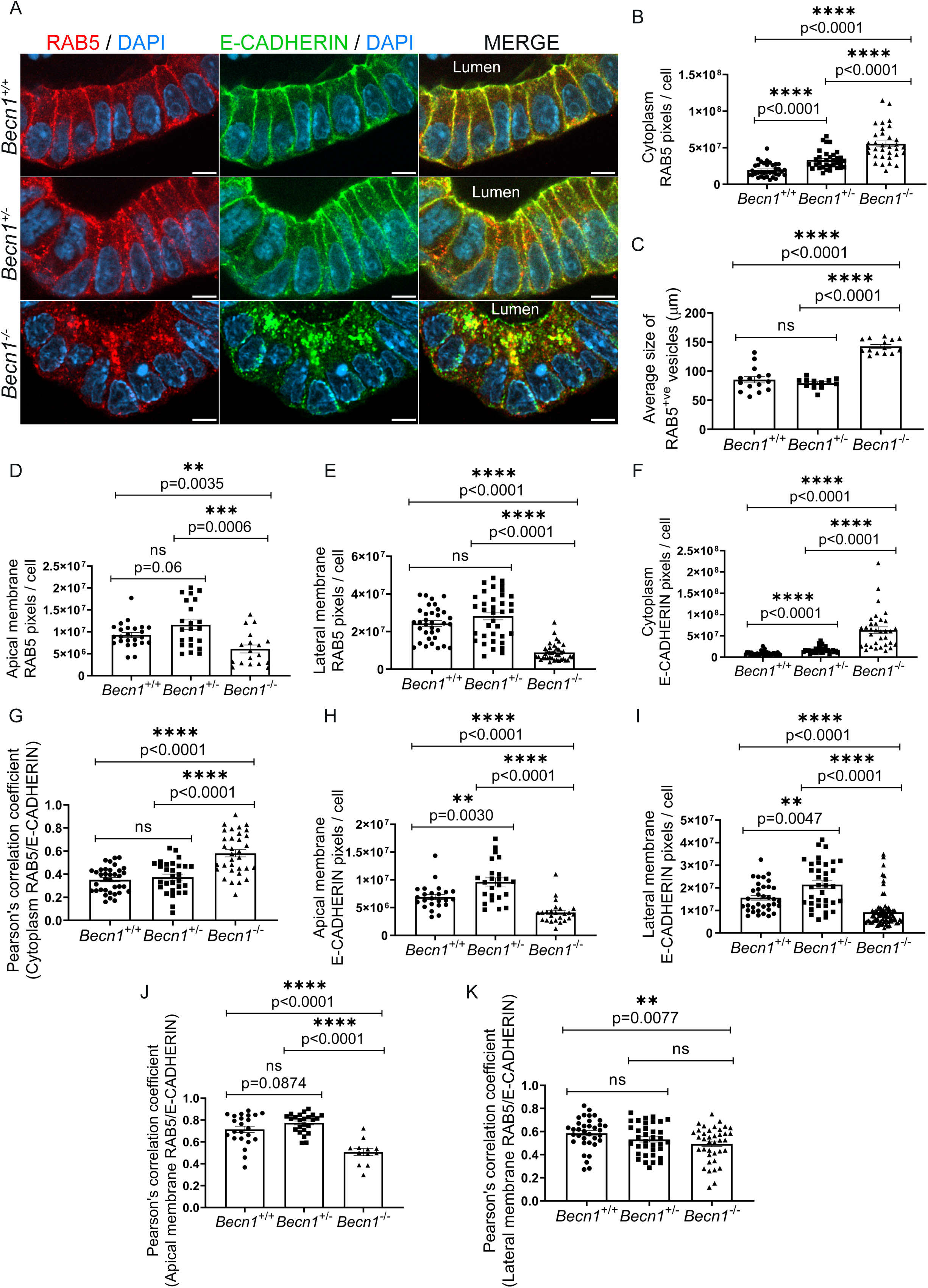
BECLIN1 reduction alters E-CADHERIN trafficking in a manner independent of RAB5^+ve^ endosomal localisation, and distinct from complete BECLIN1 deletion. **(A)** Representative whole-mount immunofluorescence staining for RAB5 (red) and E-CADHERIN (green) in *Becn1^+/+^, Becn1^+/-^ and Becn1^-/-^* organoids. Scale bar = 5 μm. **(B)** Quantification of cytoplasmic RAB5 pixels per cells. **(C)** Measurement of the average size of RAB5^+ve^ puncta. **(D)** Quantification of RAB5 signals on the apical and **(E)** lateral membranes of intestinal epithelial cells. **(F)** Quantification of cytoplasmic E-CADHERIN pixels per cells, along with its **(G)** co-localisation with cytoplasmic RAB5, as measured using Pearson’s correlation coefficient. **(H)** Quantification of apical and **(I)** lateral E-CADHERIN signals in intestinal epithelial cells. **(J)** Measurement of the degree of colocalisation between RAB5 and E-CADHERIN on the apical and **(K)** lateral membrane of intestinal epithelial cells. Data are representative of at least *n = 3* different slices per organoid and of at least *n =* 3 biologically independent organoids. For each z-section, at least *n* = 3 individual IECs with clear apical to basal delineation were used. Graphs indicate the mean ± S.E.M. Statistical significance was determined using unpaired (Student’s) t-test. ns = not significant (p > 0.05).

We next looked at the localisation of E-CADHERIN, as we previously showed that there was increased entrapment and co-localisation of this important junctional protein in the aberrantly enlarged RAB5^+ve^ early endosomes in the cytoplasm of *Becn1^-/-^*IECs (Figures 2A, F, G) ^31^. Interestingly, whilst there was increased E-CADHERIN localised at the apical and lateral membranes of *Becn1^+/-^*IECs (Figures 2H, I), as well as in the cytoplasm (Figure 2F), there were no significant changes in the amount of E-CADHERIN co-localised with RAB5 at any of these subcellular locations (Figures 2G, J, K). These results suggest that whilst reduced BECLIN1 levels can impact early endosomal dynamics, the altered localisation of E-CADHERIN may also be attributable to mechanisms independent of RAB5^+ve^ early endosome-mediated trafficking. Such mechanisms may include trafficking from the trans-Golgi network to the membrane, *via* cytoskeletal rearrangements mediated by small GTPases (e.g. RHOA, RAC1), or myosin-mediated trafficking along the actin network ^55^.

### Monoallelic loss of *Becn1^+/-^* IECs leads to changes in F-actin and cell architecture

We previously showed that homozygous loss of BECLIN1 results in cytoskeletal disruption that contributes to a compromised intestinal epithelial barrier ^30^. Therefore, to explore if changes in cytoskeletal organisation also contribute to the observed changes in E-CADHERIN localisation and trafficking in *Becn1^+/-^* organoids, we examined the distribution and dynamics of F-actin, a key structural component linked to adhesion and endocytic processes ^56, 57^. Here, there was a significant increase in the amount of F-actin and co-localised E-CADHERIN along the lateral membrane of *Becn1^+/-^* IECs that was not seen along the apical membrane (Figures 3A-E). This suggests an F-actin-driven “cadherin” flow mechanism, whereby cadherins are transported along remodelling actin networks, which could also contribute to the altered E-CADHERIN localisation observed ^58^ (Figure 2A, H, I). Indeed, E-CADHERIN is redistributed within the lateral membrane towards the apicolateral junctions, possibly as a mechanism to maintain cell-cell contacts to preserve epithelial integrity. Given the increased localisation of E-CADHERIN and F-ACTIN to the lateral membrane in *Becn1^+/-^*IECs (Figure 3E), we also examined cell length to determine whether these cell adhesion and cytoskeletal changes correlate with alterations in cell morphology, particularly whether cell elongation is impacted. We observed a significant increase in the apico-basal axis and shortening of basal IEC width in *Becn1^+/-^* organoids which likely contributes to the decreased small intestinal length in these mice (Figures 1B, 3A, F, G). These results demonstrate that reduced levels of BECLIN1 can lead to changes in adhesion dynamics and cytoskeletal organisation, as well as epithelial cell architecture to impact intestinal length.

**Figure 3.**
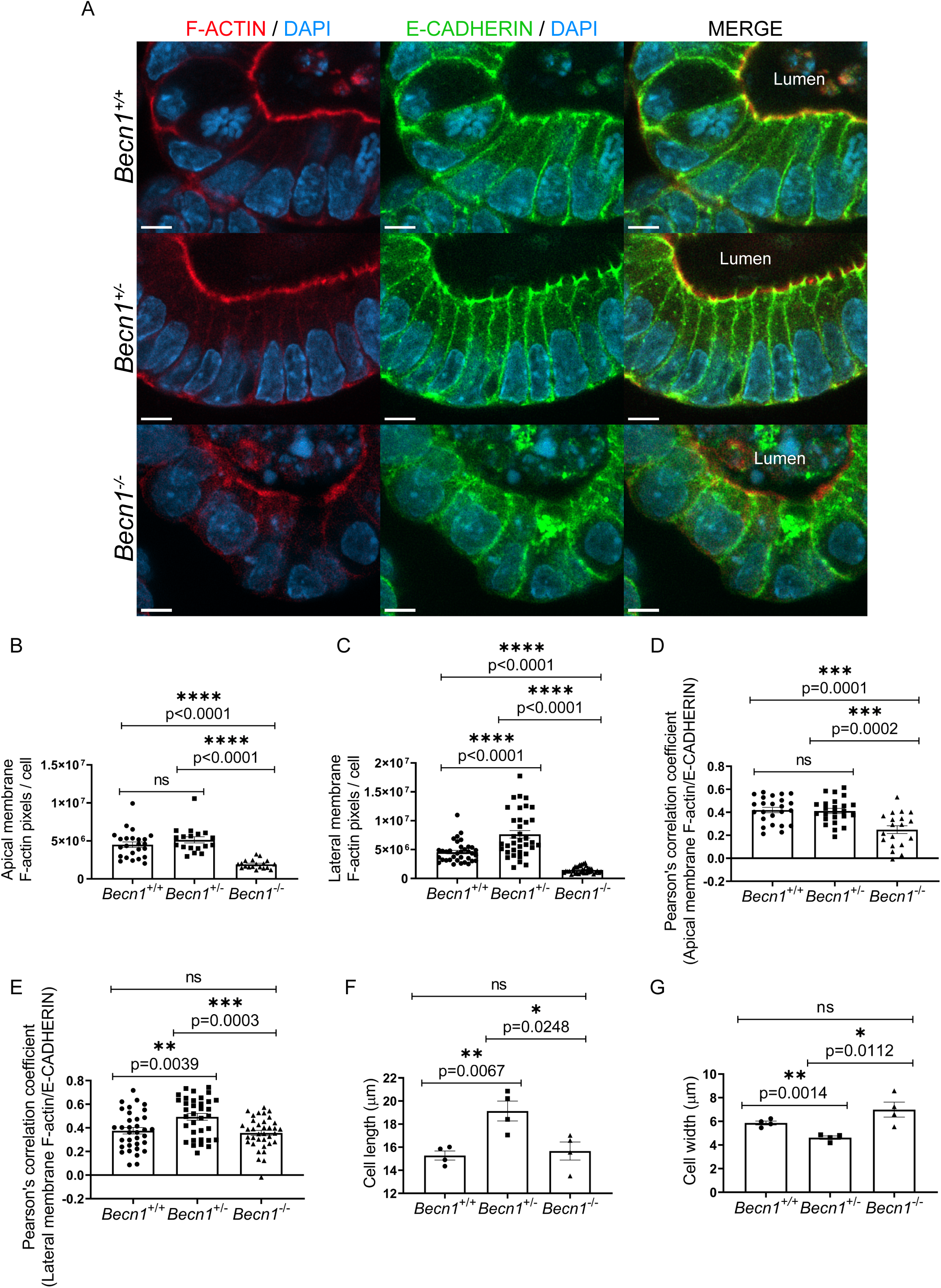
Monoallelic *Becn1* deletion alters F-actin organisation in intestinal epithelial cells. **(A)** Representative whole-mount immunofluorescence staining for F-actin (red) and E-CADHERIN (green) in *Becn1^+/+^, Becn1^+/-^ and Becn1^-/-^* organoids. Scale bar = 5 μm. **(B)** Quantification of apical and **(C)** lateral F-actin signals per cell. **(D)** Colocalisation between F-actin and E-CADHERIN on the apical and **(E)** lateral membranes of the intestinal epithelial cells, assessed using Pearson’s correlation coefficient. **(F)** Measurement of cell length (apical to the basal membrane) and **(G)** cell width (lateral membrane to lateral membrane). Data are representative of at least *n =* 3 different z-slice per organoid and of at least *n =* 3 biologically independent organoids. For each z-slice, at least *n =* 3 individual cells with clear apical to basal delineation were used for quantification. Graphs indicate the mean ± S.E.M. Statistical significance was determined using unpaired (Student’s) t-test for all graphs. ns = not significant (p > 0.05).

### Reduced levels of BECLIN1 increases susceptibility to DSS-induced colitis

Unlike *Becn1^-/-^*mice, monoallelic loss of *Becn1* in the intestinal epithelium does not lead to spontaneous fatal enteritis, disruption of gross intestinal morphology, or compromised gut barrier integrity under homeostatic conditions. Hence, we questioned if the changes in endocytic trafficking and intercellular adhesion and cytoskeleton architecture seen in *Becn1^+/-^* mice instead make them more susceptible to conditions where they are predisposed to intestinal inflammation. Accordingly, mice were treated with dextran sulphate sodium (DSS) to induce colitis, a widely used model for understanding mechanisms associated with IBD pathogenesis. For this study, mice aged 7-14 weeks with an equal distribution of sex (n=4-5 for both females and males) were used to account for sex- and developmental-related changes that can affect DSS susceptibility ^59^. Following Tamoxifen treatment to induce gene deletion, all DSS-treated mice received 2% DSS in their drinking water for 5 days, whilst control mice received normal drinking water (Figure 4A). To account for variables that could affect colitis development, such as animal weight and intake of DSS-containing water, the body weight of the mice and the weight of the DSS-containing water bottles were measured at the start and end of the DSS treatment period and did significantly different between the mice (Supplementary Figures 4A, B).

**Figure 4.**
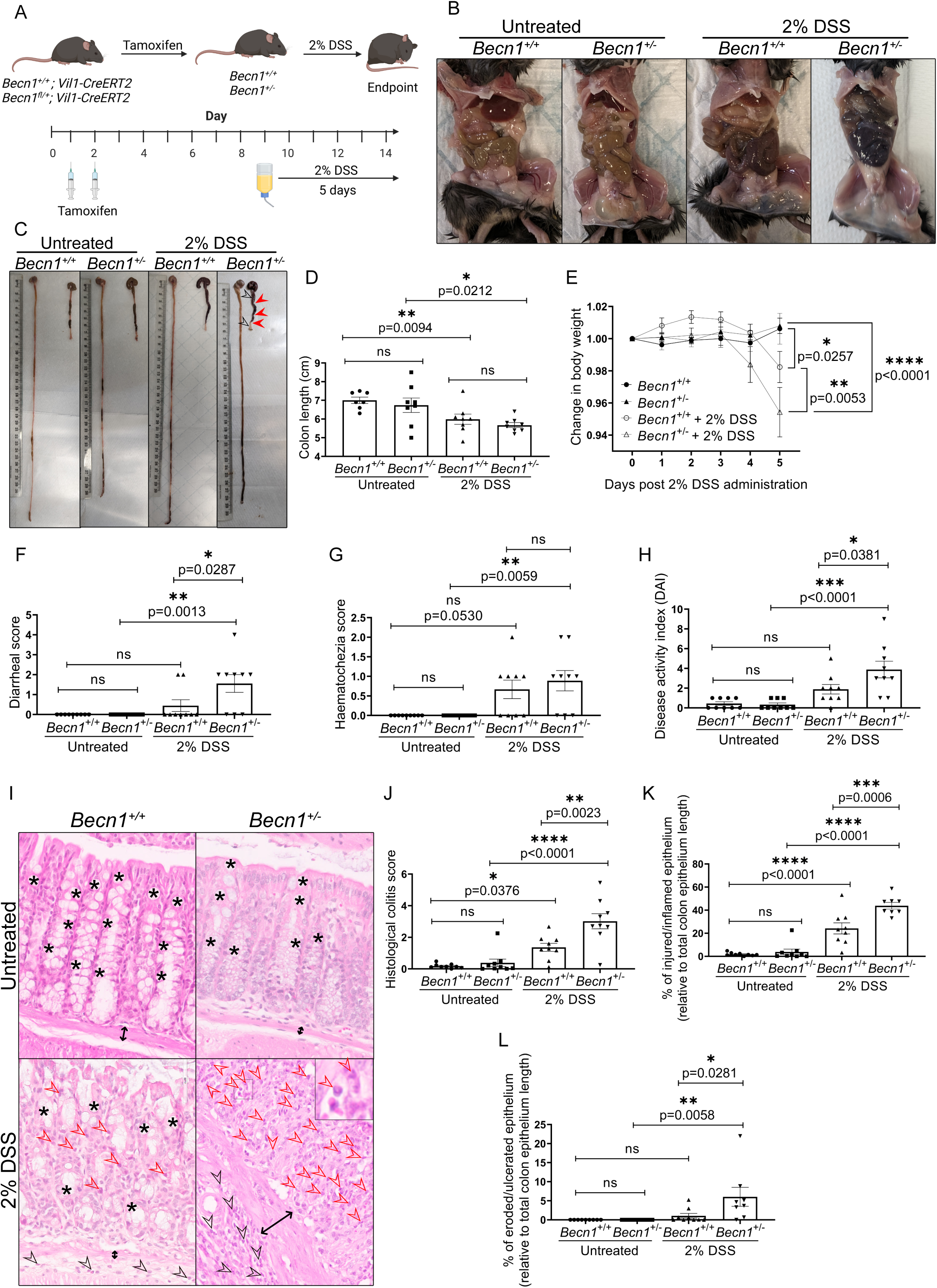
Heterozygous *Becn1* deletion exacerbates colitis severity following Dextran Sodium Sulphate administration. **(A)** Schematic diagram depicting the timeline of Tamoxifen-induced *Becn1* gene deletion followed by 5 days of 2% Dextran Sodium Sulphate (DSS) treatment. Image created in BioRender.com. **(B)** Representative images of abdominal necropsy and **(C)** intestinal tracts of *Becn1^+/+^* and *Becn1^+/-^* mice, who received normal drinking water (untreated) or 2% DSS drinking water, at endpoint. Red arrowheads and black open arrowheads highlight points of colon constriction and swelling, respectively. **(D)** Colon lengths of mice from each treatment group at endpoint. **(E)** Changes in body weight of *Becn1^+/+^* and *Becn1^+/-^* mice during the 2% DSS treatment period, normalised to body weight at the start of DSS treatment (day 9). **(F)** Diarrheal and **(G)** haematochezia scores of *Becn1^+/+^* and *Becn1^+/-^* mice who received either normal drinking water or 2% DSS drinking water. Scores were determined based on parameters outlined in Table 1. **(H)** Disease Activity Index (DAI) score of *Becn1^+/+^* and *Becn1^+/-^* mice receiving either normal drinking water or 2% DSS drinking water, calculated by summing the scores of the parameters outlined in Table 1. **(I)** HCE-stained colon FFPE sections from *Becn1^+/+^* and *Becn1^+/-^* mice who received normal drinking water (untreated) or 2% DSS drinking water (2% DSS). The colon epithelium of untreated *Becn1^+/+^* and *Becn1^+/-^* mice appeared healthy, with well-defined and uniformly distributed crypts extending from the muscularis mucosae (black double-headed arrow) to the luminal surface. The epithelial cells within the crypts are uniformly aligned with regular nuclear positioning, and goblet cells (denoted by *) are abundant. DSS-induced injury leads to crypt distortion, shortened crypts with regions of epithelial loss, triggering the recruitment of immune cells from the submucosa (black open arrowheads) into the mucosa (red open arrowhead), as shown in DSS-treated *Becn1^+/+^* mice. Notably, DSS-treated *Becn1^+/-^* mice exhibited greater susceptibility to epithelial erosion, characterised by near-total epithelial cell loss leading to complete crypt loss, loss of goblet cells and marked immune infiltration into the mucosa and submucosa compared to *Becn1^+/+^* mice. A magnified area (indicated by the box) provides a closer view of the mucosa, illustrating the increased immune infiltration in DSS-treated *Becn1^+/-^* mice. **(J)** The histological colitis score (HCS) of *Becn1^+/+^* and *Becn1^+/-^* mice with and without DSS treatment was calculated based on described methods^62^. **(K)** Percentage of injured or inflamed epithelium and **(L)** eroded epithelium, normalised to the length of the muscularis mucosae, in the colon of *Becn1^+/+^* and *Becn1^+/-^* mice receiving either normal drinking water or 2% DSS drinking water. Data are representative of at least *n =* 9 biologically independent mice from *n =* 3 independent experiments. Graphs indicate the ± S.E.M. Statistical significance was determined using ordinary one-way ANOVA except in (E) where changes in body weight was determined using two-way ANOVA with Tukey’s post-hoc test.

Gross anatomical inspection at experimental endpoint revealed expected differences between untreated and DSS-treated mice ^60^. For example, the abdominal cavities of both DSS-treated *Becn1^+/+^* IECs and *Becn1^+/-^* IECs presented with darker and reddish areas in the intestinal tract, mostly in the colon, suggestive of hyperemia and inflammation (Figures 4B, C). There was also significant shortening and contraction of the colon (Figure 4D), with noticeable areas of swelling and constriction in DSS-treated mice regardless of their genotypes (Figure 4C). At this timepoint, the small intestines of *Becn1^+/-^*mice were also significantly shorter compared to *Becn1^+/+^* controls, irrespective of whether mice received DSS treatment or not (Supplementary Figure 4C). Importantly, there were no significant differences in the extent of these gross morphological changes to the colon when comparing DSS-treated *Becn1^+/+^* to *Becn1^+/-^* mice (Figures 4B-D).

However, to conduct a more in-depth assessment of the impact of reduced BECLIN1 levels on a predisposition to inflammation, we also assessed disease severity using a composite score known as the Disease Activity Index (DAI). This scoring system evaluates parameters observed following DSS treatment including weight loss, stool consistency, and rectal bleeding (hematochezia) with the DAI score as a sum of these (Table 1) ^60, 61^. Both *Becn1^+/+^* and *Becn1^+/-^*mice demonstrated body weight loss 5 days following the commencement of DSS treatment compared to control littermates, though it was more pronounced in *Becn1^+/-^* mice (Figure 4E). Heterozygous loss of *Becn1* also resulted in more severe diarrhea following DSS administration, with more *Becn1^+/-^* mice producing pasty, semi-formed or liquid stool, as compared to *Becn1^+/+^* mice (Figure 4F). The only parameter where there was no significant difference between the genotypes was the degree of hematochezia (Figure 4G). Critically, the amalgamation of all three parameters produced a higher DAI score in the heterozygous mice, demonstrating that reduced levels of BECLIN1 increased susceptibility to DSS-induced colitis (Figure 4H).

**Table 1.** Scoring system for the evaluation of colitis severity in mice based on weight loss, stool consistency and haematochezia parameters.

| Score | Weight loss (%) | Stool consistency | Haematochezia |
| --- | --- | --- | --- |
| 0 | Normal | Normal | No blood |
| 1 | <5 |  | Visible blood in rectum |
| 2 | 6-10 | Pasty, semiformed | Visible blood on fur |
| 3 | 11-20 |  |  |
| 4 | >20 | Liquid, sticky or unable to defecate after 5 min |  |

In addition to assessment of the clinical parameters of colitis severity, we quantified the histological colitis score (HCS) of the colon following DSS treatment to evaluate the degree of inflammation and tissue damage that was not apparent with gross anatomical examination. To minimise observer bias ^59, 62^, we adopted a blinded scoring system that included assessment of the entire colon length using interpretable parameters such as crypt architecture, extent of inflammatory cell infiltration, and epithelial integrity ^62^. Unsurprisingly, minimal histological damage was observed in the colon mucosa of both *Becn1^+/+^*and *Becn1^+/-^*mice that received normal drinking water. The entire colon length presented as healthy, with organised IECs in the crypt-luminal axis and the base of the tubular glands reaching the muscularis mucosae, accompanied by few immune cells in the lamina propria (Figure 4I). These observations were reflected by the low HCS in untreated mice, with no statistical differences between the two genotypes (Figure 4J). As expected, DSS treatment led to marked increases in the HCS for both *Becn1^+/+^*and *Becn1^+/-^* mice, though this score was 2.2-fold higher in the heterozygous mice (Figure 4J). This higher HCS score for the *Becn1^+/-^* mice was attributed to a significantly more injured and inflamed colon epithelium (Figure 4K), characterised by attenuated epithelial crypts, missing epithelial cells, and/or increased immune cell infiltration (Figure 4I). There were also significantly more areas of epithelial erosion (Figure 4L) where there was a complete lack of epithelial crypts and marked localised immune infiltration (Figure 4I). Of note, there were no significant differences in lymphoid follicle count (Supplementary Figure 4D) and size (Supplementary Figure 4E) in both DSS-treated and untreated *Becn1^+/+^*and *Becn1^+/-^*mice. This suggests that the inflammation is restricted only to the epithelial layer rather than invoking systemic lymphoid tissue involvement following DSS treatment in the *Becn1^+/-^* mice.

Collectively, the increased HCS and DAI scores seen in the *Becn1^+/-^* mice compared with *Becn1^+/+^*mice following DSS administration indicate worsened colitis severity and increased sensitivity to inflammation following monoallelic *Becn1* loss (Figures 4H-L).

### Monoallelic deletion of *Becn1* leads to goblet cell defects and compromised mucosal immunity under basal conditions

Goblet cells are a secretory IEC sub-type that specialise in the production of mucus essential for the formation of a protective barrier against pathogens and mechanical damage ^63^. Disruption of the mucus barrier is a hallmark of chronic inflammatory conditions, including IBD, which often arises from goblet cell loss, or defects in mucin formation, storage, or secretion ^63–71^. A previous study showed that goblet cells in mice harbouring a constitutively active mutant of BECLIN1 had enhanced mucus secretion by alleviating ER stress through unrestrained autophagy induction ^18^. Therefore, to elucidate potential mechanisms accounting for the exacerbated colitis observed in *Becn1^+/-^* mice following DSS treatment, we investigated if there were any disruptions to the goblet cell compartment. Goblet cells in colonic tissues were histologically assessed using Periodic-Acid-Schiff (PAS) and Alcian Blue (AB) staining, which labels highly glycosylated proteins, particularly goblet cell mucins ^20^. Since DSS-induced colitis primarily affects the distal colon ^72^, we focused our analysis on this region unless otherwise stated.

Notably, untreated *Becn1^+/-^*mice displayed significantly reduced PAS-AB staining in the distal colon of *Becn1^+/-^*compared with *Becn1^+/+^*mice (Figures 5A, B). This could be attributed to decreased goblet cell numbers, maturation defects, impaired function, and/or reduced mucin production. As goblet cell maturation is associated with their migration to the colonic crypt surface, we compared the number of goblet cells in the lower (immature) and upper (mature) colonic crypt (Figure 5C, Supplementary Figure 5) to assess the impact of reduced BECLIN1 levels on maturation ^20, 66, 73^. There was reduced PAS-AB^+ve^ mucin staining in both halves of the colonic crypt in *Becn1^+/-^* mice though this was more pronounced in the upper halves (50% reduction compared to 15% reduction in the lower halves) (Figure 5C). This apparent maturation defect is further underscored by the larger cytoplasmic mucin, or theca, areas seen in the remaining goblet cells in the upper crypts of *Becn1^+/-^* mice compared with *Becn1^+/+^*mice, indicative of impaired mucin secretion (Figure 5D). Hence, these results indicate reduced BECLIN1 is associated with both an overall loss of goblet cells and/or defects with mucin production alongside defective goblet cell maturation and migration from the crypt base to the luminal end.

**Figure 5.**
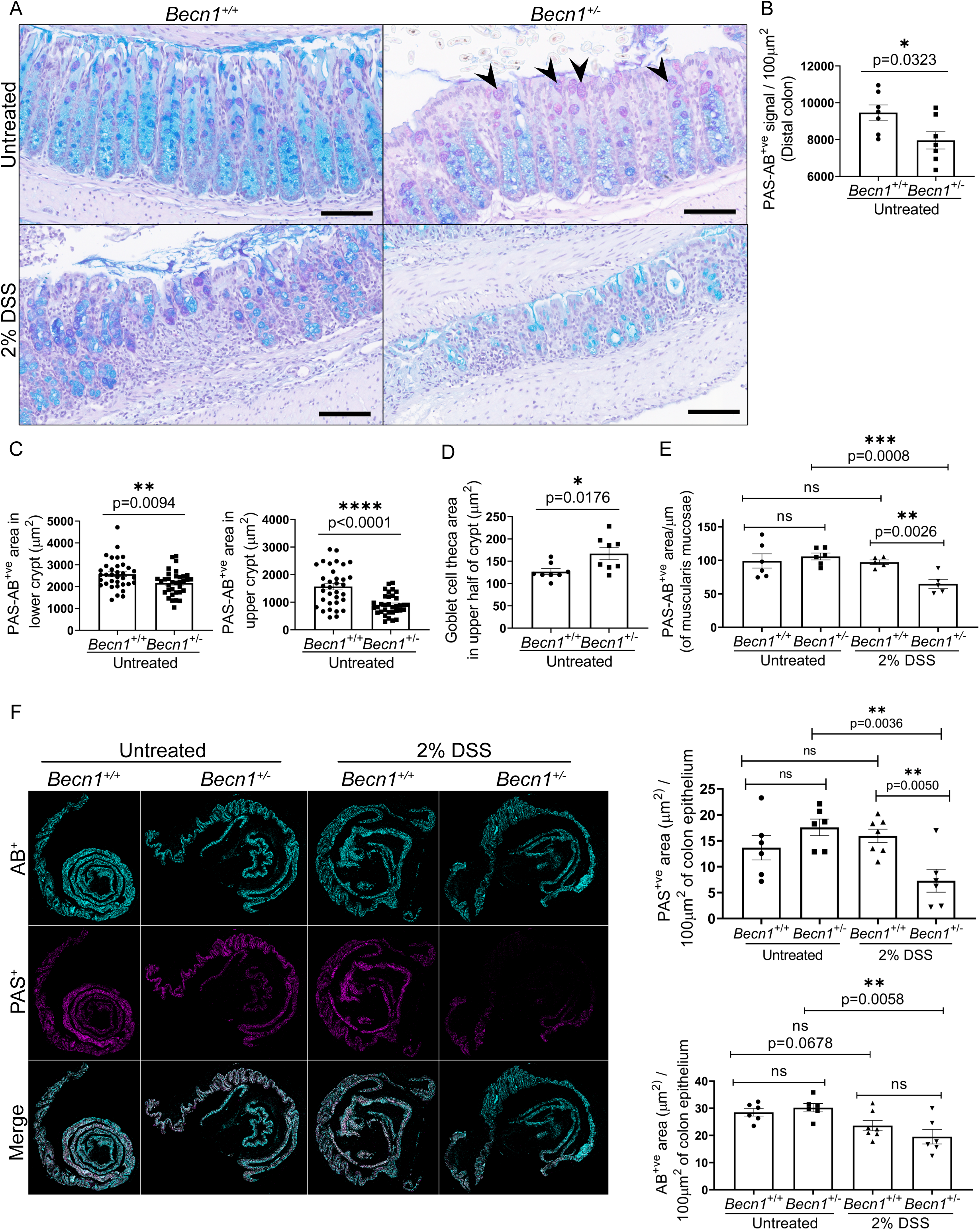
Heterozygous *Becn1* deletion disrupts goblet cell function and predisposes to DSS-induced colitis. **(A)** Representative PAS-AB-stained distal colon sections from untreated and DSS-treated *Becn1^+/+^* and *Becn1^+/-^*mice. Scale bar = 100 μm. Black arrowheads depicts enlarged goblet cells. **(B)** Quantification of the number of PAS-AB^+ve^ signal in distal colon epithelium of *Becn1^+/+^* and *Becn1^+/-^* mice. **(C)** Quantification of PAS-AB^+ve^ area in the top half and bottom half of distal colon crypts, as delineated in Supplementary Figure 5, in *Becn1^+/+^* and *Becn1^+/-^* mice. **(D)** Quantification of goblet cell theca size in the top half of distal colonic crypts of *Becn1^+/+^*and *Becn1^+/-^*mice. **(E)** Quantification of PAS-AB^+ve^ area relative to total colon length (measured relative to muscularis mucosae length) in untreated and DSS-treated *Becn1^+/+^* and *Becn1^+/-^* mice. **(E)** Graphical representation of acidic (AB^+ve^), neutral (PAS^+ve^), and total (Merge) mucins in untreated and DSS-treated *Becn1^+/+^* and *Becn1^+/-^* mice. The area of acidic (AB^+ve^) and neutral (PAS^+ve^) mucus were also quantified per 100 μm^2^ of colon epithelium. Data represent n > 5 biologically independent mice from *n =* 2 independent experiments. Only animals that received DSS treatment from the same lot number were included for analysis. Graphs show the ± S.E.M. Statistical significance was determined by unpaired (Student’s) t-test. For crypt-specific quantifications in (C), full length intact crypts with at least three contiguous intact crypts were analysed, with *n >* 3 crypts analysed per animal. The average goblet cell theca area in (D) was obtained by measuring >100 goblet cells per animal. DSS: dextran sodium sulphate. PAS: periodic acid Schiff. AB: alcian blue. ns: not significant (p > 0.05).

### Reduction of BECLIN1 exacerbates DSS-induced epithelial damage and mucosal barrier dysfunction

Despite PAS-AB staining being significantly reduced in the distal colon of untreated *Becn1^+/-^* mice (Figures 5A, B), PAS-AB staining when measured across the total colonic epithelial surface (i.e. including both the proximal and distal colon) revealed no significant differences between untreated *Becn1^+/+^* and *Becn1^+/-^* mice (Figure 5E). This suggests an increase in mucin production in the proximal colon under homeostatic conditions to compensate for the decreased mucin in the distal colon of *Becn1^+/-^* mice. We next investigated goblet cell responses in DSS-treated *Becn1^+/+^* and *Becn1^+/-^*mice. Similar to under basal conditions, DSS-treated *Becn1^+/+^* mice showed no significant changes in total PAS-AB^+ve^ mucin area in the entire colon compared to untreated *Becn1^+/+^* controls, despite significant evidence of epithelial damage in the distal colon (Figure 4I-L). This indicates once again a likely compensatory increase in mucin production in the proximal colon in response to DSS-induced injury. In contrast, DSS treatment of *Becn1^+/-^* mice resulted in a reduction of approximately two-fold, in the total colonic PAS-AB^+ve^ area compared to DSS-treated *Becn1^+/+^*, and to untreated *Becn1*^+/-^ mice (Figure 5E). Specifically, we observed a significant loss of PAS^+ve^ neutral mucins, but not AB^+ve^ acidic mucins, in the colons of DSS-treated *Becn1^+/-^*mice compared with wild-type mice (Figure 5F). This disproportionate loss of PAS^+ve^ neutral mucins highlights a specific vulnerability in the mucosal barrier when BECLIN1 levels are reduced reinforcing a role for BECLIN1 in maintaining epithelial homeostasis and underscoring the importance of neutral mucins in protecting against DSS-induced colitis.

### Reduced levels of BECLIN1 are associated with endoscopically inflamed regions of intestinal biopsies from patients with IBD

Combined, the above results demonstrate that reduced levels of BECLIN1 lead to increased susceptibility to DSS-induced colitis, which is a widely used model of IBD, specifically mirroring the characteristics of human ulcerative colitis. We next wanted to translate these *in vivo* findings from our mouse model into a more clinically relevant context by examining if BECLIN1 levels are reduced in intestinal biopsies from IBD patients. Western blotting was performed on biopsies collected from endoscopically “non-inflamed”, “marginally inflamed”, and “inflamed” intestinal tissue from IBD patients ^74, 75^. BECLIN1 levels in the lysate from non-IBD patients was used as a comparator. We observed a general trend towards increased levels of BECLIN1 in marginally inflamed regions when compared to the non-IBD and non-inflammed tissue in the majority of the patient samples (Figure 6). Notably, whilst there was heterogeneity between the samples analysed, BECLIN1 levels were greatly reduced in the inflamed regions of three out of the five IBD patient samples (Figure 6). These early clinical data demonstrates that BECLIN1 levels can change regionally in IBD pathogenesis and that decreased BECLIN1 can be associated with inflammation of the intestinal epithelium. This is consistent with our *in vivo* findings where monoallelic deletion of BECLIN1 in the intestinal epithelium of mice leads to increased susceptibility to DSS-induced colitis.

**Figure 6.**
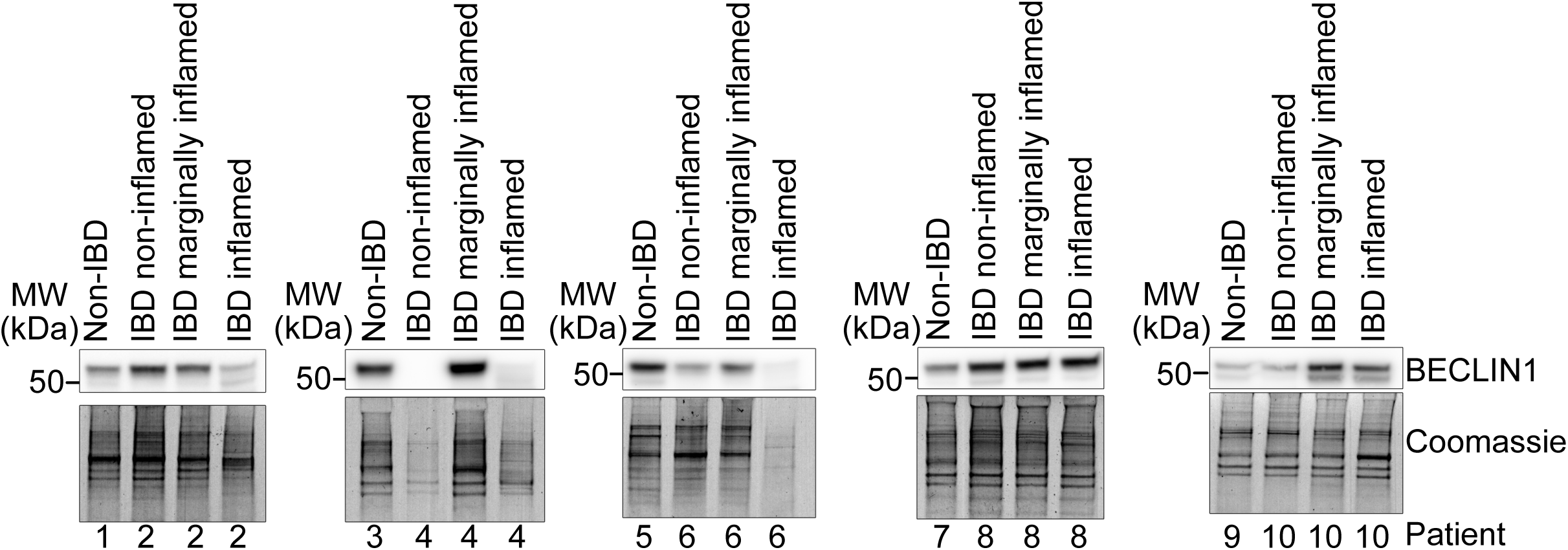
Analysis of BECLIN1 expression in IBD patient samples. Western blot analysis of tissue biopsies isolated from inflammatory bowel disease (IBD) patients (2, 4, 6, 8, and 10) along with non-IBD patient (1, 3, 5, 7 and 9) as reference. Coomassie stained gels were used as loading controls.

## DISCUSSION

We previously identified an essential role for BECLIN1 in maintaining intestinal homeostasis and barrier integrity mediated through its autophagy and endocytic trafficking functions ^30, 31^. Given the phenotypic resemblance of BECLIN1 homozygous knockout mice to some of the clinical manifestations of IBD, we sought to investigate the potential contributions of reduced BECLIN1 levels (i.e. a more disease relevant setting) to intestinal homeostasis, susceptibility to inflammation, and in turn IBD pathogenesis.

Unlike homozygous BECLIN1 deletion, heterozygous deletion did not lead to spontaneous and fatal intestinal disruptions in adult mice ^31^. Instead, heterozygous BECLIN1 loss compromised the structural integrity of colonic crypts leading to a significant shortening of crypt lengths in the distal colon, a phenomenon indicative of crypt atrophy. Crypt atrophy can be attributed to several factors, including reduced colonic stem cell proliferation, or increased cell death resulting in the loss of functional epithelial cells such as absorptive or Goblet cells ^76^. Our data suggest the shortened crypt lengths in the *Becn1^+/-^* colon is unlikely due to compromised proliferation but due to a loss of goblet cells. This would be consistent with the overall reduction in goblet cell numbers along the entire crypt length and the loss of this cellular compartment in homozygous BECLIN1 knockout mice ^31^. In addition, altered F-actin localisation in IECs, such as that seen in *Becn1^+/-^* cells, has been linked to disrupted IEC contractility and morphological changes that affect crypt architecture ^77^. Crypt atrophy can be an indicator of predisposition to gastrointestinal pathologies such as IBD by influencing colonic function due to changes in immune responses or nutrient absorption and has been documented in conditions such as ulcerative colitis ^49, 78–81^.

In addition to the manifestation of crypt atrophy, reduced BECLIN1 levels also led to maturation and mucin abnormalities in goblet cells. These were evidenced by a more pronounced decrease in mucin-positive cell numbers, and an increase in theca size observed in remaining goblet cells located in the upper half, as compared to those in the lower half, of the colonic crypt in untreated mice. Theca size correlates with the number of stored mucin granules within goblet cells, and an increase in size can be attributed to several factors including a compensatory response to loss of surrounding goblet cells or the abnormal retention of mucin granules through defective exocytosis, or impaired maturation of goblet cells ^14, 20, 70^. The regulation of mucin secretion by goblet cells is partially mediated by F-actin filaments, which negatively regulate secretion by restricting mucin granule access to the plasma membrane for exocytosis ^82^. Disruption of F-actin accelerates mucin granule movement enhancing baseline mucin secretion, whilst its stabilisation inhibits mucin release ^82, 83^. The enrichment of laterally localised F-actin on the *Becn1^+/-^* IEC membrane, and its increased co-localisation with E-CADHERIN, likely stabilises cell-cell adhesion and the F-actin cytoskeleton ^84^. It is therefore possible that this stabilisation inhibits mucin secretion as seen in *Becn1^+/-^*IECs. However, this hypothesis requires further validation. Notably, these secretory defects align with those observed in homozygous BECLIN1 knockout mice, where we previously identified defects in other secretory IECs, such as Paneth cells, displaying swollen lysozyme granules and an increased proportion of depleted cells containing fewer granules with reduced lysozyme levels ^31^.

Proper goblet and Paneth cell function clearly relies not only on autophagy but also on intact endocytic trafficking. Studies of autophagy-related genes such as *Atg5*, *Atg1CL*, and trafficking regulators like RAB7 highlight how disruptions in these pathways impair mucin granule handling and mucus secretion, ultimately sensitising the gut to inflammation ^14, 20, 85^. Previously, BECLIN1 has been shown to facilitate the proper secretion of mucus from colonic goblet cells through its role in the autophagy-mediated suppression of ER stress ^18^. This present study indicates that its autophagy-independent roles also contribute to goblet cell function. Hence, these collective observations underscore the notion that reduced BECLIN1 levels contribute to a predisposition to inflammatory intestinal pathologies, such as IBD, through its maintenance of goblet cell function. Whilst our data show that heterozygous BECLIN1 knockout mice do not develop spontaneous gastrointestinal pathologies within the short time frame of this study, aging or further exposure of these mice to additional stressors could trigger the development of such indications mediated by goblet cell dysfunction.

Indeed, mice heterozygous for *Becn1* exhibited heightened susceptibility to DSS-induced colitis, accompanied by a shift in goblet cell mucin composition toward a more acidic profile due to a marked reduction in neutral mucins. While acidic mucins enhance mucus viscosity and resistance to bacterial degradation, neutral mucins are important for buffering luminal acidity and neutralising toxic compounds. Similar shifts have been reported in IBD patients and may reflect an adaptive response to inflammation^86–91^. However, in *Becn1*^⁺/⁻^ mice, this response was insufficient to compensate for the overall loss of mucins, impaired goblet cell function, and compromised barrier integrity, contributing to increased epithelial damage and inflammation.

Our study demonstrates that maintaining appropriate BECLIN1 levels is critical for mucin secretion and preserving the mucus barrier, a key defence against intestinal inflammation. Disruption of this barrier, often due to goblet cell loss or mucin defects, is a hallmark of chronic inflammatory diseases such as IBD ^63–71, 92, 93^. Consistent with our in vitro and animal findings, analysis of IBD patient biopsies revealed that BECLIN1 expression varied with inflammation status. Levels of BECLIN1 tended to rise in marginally inflamed regions and decline markedly in areas of severe inflammation as compared to non-inflamed or non-IBD samples. This spatial variation may explain previous discrepancies in the literature regarding BECLIN1 expression in IBD^47, 94–97^. Our findings thus provide novel spatial context to BECLIN1 regulation in human disease, suggesting dynamic modulation during disease progression.

The question that arises is whether these changes in BECLIN1 levels are the cause, or the effect, of increased susceptibility to intestinal inflammation. If the former, these observations suggest a causative cytoprotective role for BECLIN1 in mitigating the detrimental effects of inflammation and/or apoptosis inhibition. Previous studies have shown a role for BECLIN1-mediated autophagy in suppression of ROS accumulation, the autophagic degradation of danger signals that activate the inflammasome, or by alleviating ER stress in IECs leading to increased mucus barrier protection ^18, 20, 98^. There is also precedence for autophagy-independent mechanisms by which BECLIN1 protects against inflammation^99–104^. Interestingly, whilst no IBD-associated SNPs have been identified within BECLIN1 in GWAS analyses, our data also suggest that it may be pertinent to investigate upstream regulators of BECLIN1 expression and activity instead. For instance, SNPs associated with increased expression and/or activation of STAT3, a transcription factor that has been shown to transcriptionally repress *Becn1* expression through the recruitment of HDAC3 to the *Becn1* promoter, have been implicated in IBD pathogenesis ^105–107^. This raises the possibility that alterations in transcription factor signalling could indirectly influence BECLIN1 levels, contributing to the compromised epithelial and mucosal barrier function observed in IBD.

Alternatively, the reduced BECLIN1 levels associated with highly inflamed intestinal biopsies could be due to consequential mechanisms such as caspase-mediated cleavage as previously reported ^108, 109^. Such a mechanism would be expected to create an amplifying loop enhancing the apoptotic signal. Given that caspase-mediated cleavage of BECLIN1 has been described^108, 109^, we propose that one plausible explanation for the heterogeneity in BECLIN1 levels observed in inflamed tissues across patient samples may be a reflection of the mode of cell death signalling occurring in the intestinal epithelium ^110^. Expanding the clinical cohort will enable us to validate these changes in BECLIN1 levels across a broader population and draw statistically significant conclusions strengthening the clinical relevance of our findings.

Our results pave the way for future work to validate BECLIN1 as a biomarker for IBD pathogenesis and disease severity, with the potential to stratify patients and guide therapeutic interventions. Hence, this work emphasises the importance of BECLIN1 not only in epithelial homeostasis but also identifies a promising avenue for advancing precision therapeutic intervention in IBD management.

## MATERIALS AND METHODS

### Mice

All mouse strains used in this study were bred on the C57BL/6 J background. Becn1^tm1b(KOMP)Wtsi^ mice were purchased from the European Conditional Mouse Mutagenesis Program (EUCOMM). Becn1^fl/fl^ mice were generated by breeding Becn1tm1b(KOMP)Wtsi mice onto CAG-FLPe mice. *Becn1^+/+^;*, *Becn1^+/ff^;,* and *Becn1^ff/ff^*; *Vil1-CreERT2^Cre/+^* mice were then subsequently generated by breeding *Becn1^+/+^*, *Becn1^fl/+^* and *Becn1^ff/ff^* mice to the Vil1-CreERT2 mice^111^.

Mice were housed at the La Trobe Animal Research and Teaching Facility (LARTF, La Trobe University, VIC, Australia) under Specific Pathogen Free (SPF) conditions. All experiments performed were approved by the La Trobe University animal ethics committees (approvals AEC18024, AEC18036) in accordance with the Australian code for the care and use of animals for scientific purposes. All research with these mice has complied with all relevant ethical regulations for animal use. To induce deletion, male and female mice selected indiscriminately, aged six weeks or older, were intraperitoneally injected with 4 mg tamoxifen (Sigma-Aldrich, T5648) in sunflower seed oil (Sigma-Aldrich, 25007), delivered as one 200 μl injection per day of a 10 mg/ml stock, over two consecutive days. Mice were humanely end-pointed by CO_2_ asphyxiation.

### Dextran Sodium Sulphate treatment

Following Tamoxifen treatment to induce gene deletion, mice were administered DSS (2% (w/v), MP Biomedicals, Canada) in autoclaved drinking water provided ad libitum for 5 days. Control groups received autoclaved drinking water without DSS for the same duration. During the experimental period, mice were monitored daily and weights taken. The average daily intake of 2% (w/v) DSS water was calculated by measuring the weight difference of drinking bottles at the start and end of the experiment and dividing by the number of treatment days. The disease severity score was assessed using a cumulative scoring system based on parameters outlined in Table 1, including weight loss, stool consistency and the presence of hematochezia (presence of blood in stool). At the experimental endpoint, mice were humanely euthanised using CO_2_ asphyxiation. The small and large intestine were harvested, and their lengths were measured prior to Swiss-rolling and fixation in 10% (v/v) neutral buffered formalin for 48 hours.

### PCR genotyping

For colony maintenance and gene deletion assessments, DNA was extracted from ear clips or isolated IECs (see IEC isolation method). Samples were incubated overnight at 56°C in DirectPCR (Tail) Lysis Reagent (Viagen) supplemented with 0.2% (v/v) Proteinase K (Sigma-Aldrich), followed by heat inactivation at 85 °C for 1 hour. PCR amplification was performed using GoTaq PCR Master Mix (Promega) according to the manufacturer’s protocol. Primers for *Becn1* reactions are as follows: “floxed forward”—5′CTG ATC CTG CAG CTT GCA GAT TAG C3′, “floxed reverse”—5′CAC CAC TGC CTG GCT AAA CAA GAG C3′, “KO reverse”—5′CTA TAG AAG AAA GGA CTG TTG TGA AC3′. Primers for *Vil1-CreERT2* reactions are as follows: forward—5′CAA GCC TGG CTC GAC GGC C3′, reverse - 5′CGC GAA CAT CTT CAG GTT CT3′. All primers were purchased from Integrated DNA Technologies as single stranded sequences purified by standard desalting and were resuspended at 10 OD/ml in nuclease-free H2O as working stock solutions. Reactions were cycled as follow: *Becn1* (Step 1: 94° C for 4 minutes, Step 2: 94° C for 30 seconds, Step 3: 55° C for 30 seconds, Step 4: 72° C for 1 minute, Step 5: Repeat steps 2-4 for an additional 29 times, Step 6: 72° C for 5 minutes, Step 7: Hold at 4° C) and *Vil1-CreERT2* (Step 1: 95° C for 5 minutes, Step 2: 94° C for 45 seconds, Step 3: 62° C for 25 seconds, Step 4: 72° C for 1.15 minute, Step 5: Repeat steps 2-4 for an additional 34 times, Step 6: 72° C for 5 minutes, Step 7: Hold at 4° C). Reaction products were electrophoresed on 1.5% (w/v) agarose gels containing SYBR^TM^ Safe DNA Gel Stain (Invitrogen^TM^) diluted at 20,000x in TAE buffer (40mM Tris Base, 20mM glacial acetic acid, 1mM EDTA). In all gels, 1 Kb Plus DNA Ladder (Invitrogen^TM^) was used as a marker for DNA migration according to size. DNA was visualised *via* UV excitation using the ChemiDoc^TM^ Imaging System (Bio-Rad) and images processed with Image Lab (Bio-Rad).

### Intestinal organoid culture

Organoids were established by culturing crypt-enriched fractions from the duodenum of untreated mice as described previously ^31^. The crypts were cultured in Cultrex Reduced Growth Factor Basement Membrane Extract, Type 2, Path Clear (RCD Systems) at a density of 50-300 crypts per 50 μl dome and plated onto pre-warmed 24 well plates. Domes were allowed to polymerise by incubation at 37 °C for 15 minutes and Advanced DMEM media supplemented with 1× N-2 (Gibco), 1× B-27 (Gibco), 0.05 ng/μl human EGF (Peprotech, AF-100-15), 0.1 ng/μl murine noggin (Peprotech, 250-38), 0.5 ng/μl murine R-Spondin-1 (Peprotech, 315-32). Organoids were maintained at 37°C, 5% CO_2_, with media changes performed every 2-3 days. Passaging was done every 7-10 days by mechanical dissociation and re-seeding into fresh Cultrex at a 1:3 split ratio. To induce *Becn1* deletion, organoids were seeded into media containing 200 nM 4-hydroxytamoxifen (4-HT) (Sigma-Aldrich, H7904) for three days, and then maintained as per normal. For organoid re-passaging experiment, 4-HT-treated organoids were passaged at day 7 and maintained as above.

### Intestinal epithelial cell isolation

Intestinal sections were dissected open longitudinally, rinsed with dPBS, and incubated in 15 mM EDTA prepared in DPBS for 15 minutes at 37 °C with agitation. Dissociation of IECs was achieved by vortexing. The cell suspension was then washed once with ice-cold dPBS, and the cell pellet snap-frozen on dry ice and stored at −80 °C until further use.

### Immunoblotting

Cells were lysed on ice for 1 hour in lysis buffer containing 20 mM Tris pH 7.5, 135 mM NaCl, 1.5 mM MgCl_2_, 1 mM EDTA, 10% (v/v) glycerol, 1% (v/v) Triton X-100, supplemented with cOmplete Protease Inhibitor Cocktail (Roche, following the manufacturer’s protocol). Lysates were centrifuged at 16,000 × *g* for 5 minutes and supernatant protein concentrations determined using the Pierce BCA Protein Assay Kit (Thermo Scientific) according to the manufacturer’s protocol. Equalised protein amounts (15-40 µg per lane) were boiled for 5 minutes at 95 °C in reducing buffer (4× stock prepared at 178.3 mM Tris, 350 mM dithiothreitol, 288.4 mM SDS, 670 mM bromophenol blue, 36% (v/v) glycerol, 10% (v/v) β-mercaptoethanol) and electrophoresed on 4– 12% polyacrylamide NuPAGE™, Bis-Tris Mini Protein Gels (Invitrogen) using NuPAGE™ MES SDS Running Buffer (Invitrogen). Proteins were transferred onto 0.45 μm nitrocellulose membranes (Amersham Protran) *via* wet transfer at 17 V for 1 hour in in NuPAGE™ Transfer Buffer supplemented with 10% (v/v) methanol, using the Mini Blot Module (Invitrogen). Membranes were blocked in 5% (w/v) skim milk in dPBS (2.7 mM KCl, 1.76 mM KH_2_PO_4_, 136.7 mM NaCl, 8.07 mM anhydrous Na_2_HPO_4_) for 1 hour at RT with agitation. Primary antibody incubations were performed overnight at 4 °C in 1% (w/v) skim milk in PBST (PBS + 0.05% (v/v) Tween-20) with the following dilutions: Beclin1, 1:500 (CST, 3495); p62/SQSTM1, 1:500 (CST, 5114); LC3B, 1:500 (Novus Biologicals, NB100-2220); β-Actin, 1:5000 (Sigma-Aldrich, A2228); GAPDH, 1:5000 (Invitrogen, MA5-15738). Membranes were washed in PBST and probed for 1 hour at RT with the following secondary antibodies, prepared in 1% (w/v) skim milk in PBST: Donkey anti-Rabbit IgG, 1:10,000 (GE Healthcare, NA943V); Goat anti-Mouse IgG, 1:10,000 (Sigma-Aldrich, A0168). Luminescent signals were visualized using the Western Lightning Plus-ECL (PerkinElmer) kit and ChemiDoc Imaging System (Bio-Rad). Images were processed using Image Lab (Bio-Rad).

For human intestinal IBD biopsy analysis, lysates were boiled for 10 minutes in Laemmli sample buffer (126 mM Tris-HCl, pH 8.0, 20% (v/v) glycerol, 4% (w/v) SDS, 0.02% (w/v) bromophenol blue, 5% (v/v) 2-mercaptoethanol) and separated by SDS-PAGE on 4–12% Bis-Tris gels (Thermo Fisher Scientific, NP0335BOX) using MES running buffer (Thermo Fisher Scientific, NP000202). Proteins were transferred onto polyvinylidene difluoride (PVDF) membranes (Merck, IPVH00010), and gels were Coomassie-stained following the manufacturer’s protocol (Thermo Fisher Scientific, LC6060). Membranes were blocked in 5% (w/v) skim milk powder in TBS containing 0.1% (v/v) Tween-20 (TBS-T).

Primary antibody incubation was performed overnight at 4°C in blocking buffer containing anti-Beclin1 antibody (1:1000; CST, 3495). Membranes were washed twice in TBS-T and incubated with horseradish peroxidase (HRP)-conjugated secondary antibodies at 1:10,000 dilution of anti-rabbit IgG (Southern Biotech, 4010-05). Following four washes in TBS-T, signals were detected using enhanced chemiluminescence (Merck, WBLUF0100) and imaged with the ChemiDoc Touch Imaging System (Bio-Rad).

### Intestinal organoid whole-mount immunofluorescence

Whole-mount staining of intestinal organoids was performed as previously described, based on an adapted protocol from Dekkers *et al*. 2019 with modifications to the clearing and mounting steps ^112^. Briefly, organoids were removed from Cultrex by incubation with ice-cold Gentle Cell Dissociation Reagent (STEMCELL Technologies,100-0485) with gentle rocking at 4°C for 60 minutes. Organoids were subsequently fixed in 4% (w/v) paraformaldehyde solution (ProSciTech, C004) and blocked with Organoid Wash Buffer (dPBS (Gibco^TM^, 14190144) + 0.1% (w/v) TritonX-100 (Merck) + 0.2% (w/v) Bovine Serum Albumin (Merck, A3059)). Samples were then incubated overnight at 4°C with gentle agitation using primary antibodies at the following dilutions: E-CADHERIN, 1:500 (Invitrogen, 13-1900); RAB5, 1:100 (CST, 46449); F-actin, using 1X Phalloidin-iFluor 647 solution (Abcam, ab176759). Following primary antibody incubation, organoids were extensively washed and incubated overnight at 4°C with the following secondary antibodies or probes with gentle agitation: Goat anti-Rat IgG (H+L) cross-adsorbed secondary antibody, Alexa Fluor^TM^ 568 (1:400, Invitrogen, A-11077); Goat anti-Mouse IgG (H+L) cross-adsorbed secondary antibody, Alexa Fluor^TM^ 488 (1:400, Invitrogen, A-11004); and nuclear stain DAPI (1 μg/ml, Merck, D9542). Organoids were then subjected to another extensive washing step prior to sample clearing and mounting steps. Imaging was performed using a Zeiss LSM 980 confocal microscope equipped with Airyscan2. The FastAiryscan 2 SR-4Y mode was used, and imaging parameters such as laser power and gain were kept consistent for each repeat. Images were captured using either the 20x or 40x (water) objective with 1.7x Zoom and imaged once to avoid excessive photobleaching. Z-stacks were acquired for each image at 5 μm intervals.

### Histology

Intestinal Swiss-rolls were fixed in 10% (v/v) neutral buffered formalin for 48 hours and subsequently transferred to 80% (v/v) ethanol. The samples were sent to the Anatomical Pathology department (Austin Pathology, VIC, Australia) for processing and embedding into paraffin blocks. Sections were cut to a thickness of 4 μm and mounted onto SuperFrost Plus^TM^ (Thermo Scientific^TM^) positively charged glass slides.

The sections were dewaxed with xylene (two 5-minute incubations) and rehydrated in reverse ethanol gradients: two 2-minute incubations in 100% ethanol, one 2-minute incubation in 70% (v/v) ethanol and a final rinse in running RO water for 2 minutes. Following rehydration, the sections were stained using specific protocols outlined below. Once stained, the slides were dehydrated through ascending ethanol concentrations (one 2-minute incubation in 70% (v/v) ethanol, followed by two 2-minute incubations in 100% ethanol), cleared in xylene (two 5-minute incubations) and mounted using DPX (Sigma-Aldrich, 06522). All slides were scanned using the Aperio AT2 (Leica) and images captured and analysed using Aperio ImageScope v12.4.2.5010 and Fiji ImageJ.

### Haematoxylin and eosin (HGE) staining

Following standard deparaffinisation and rehydration, the slides were incubated in Mayer’s haematoxylin (Amber Scientific, MH) for 30 seconds to 3 minutes, followed by a 3-minute rinse under warm running tap water. The slides were then immersed in Scott’s solution (Amber Scientific, SCOT) for the same duration as the haematoxylin incubation to achieve bluing and subsequently rinsed in tap water for 1 minute. Next, the slides were incubated in 1% (v/v) aqueous eosin (Amber Scientific, EOA1), rinsed in tap water until the water ran clear, and then dehydrated, cleared and mounted as per above.

### Periodic acid-Schiff, Alcian blue (PAS-AB) staining

Tissue sections were subjected to standard deparaffinisation and rehydration steps outlined above, followed by a 15-minute incubation in 1% (v/v) Alcian blue prepared in 3%(v/v) aqueous acetic acid (Amber Scientific, ALCB). Slides were subsequently rinsed under running tap water for 2 minutes and further rinsed in RO water for an additional 2 minutes. Periodic acid oxidation was carried out by incubating the sections in 1% (v/v) aqueous period acid (Amber Scientific, PERO) for 5 minutes, followed by a 2-minute rinse in RO water. Slides were then immersed in Schiff’s reagent (Amber Scientific, SCHF) for 10 minutes and thoroughly rinsed under running tap water for 5 minutes to remove excess reagent.

Counterstaining was performed using Mayer’s haematoxylin for 30 seconds to 3 minutes, followed by a 2-minute rinse in tap water. Slides were subsequently blued in Scott’s solution for a duration equivalent to the haematoxylin incubation and rinsed again in tap water for 2 minutes. After staining, the slides underwent standard dehydration, clearing and mounting procedures outlined above.

### Immunohistochemistry (Ki67 staining)

Following standard deparaffinisation and rehydration steps described above (see Histology), antigen retrieval was performed by boiling slides in 10 mM sodium citrate with 0.05% (v/v) Tween-20, *via* boiling in the microwave for 20 minutes. Slides were cooled for 20 minutes at room temperature and rinsed with RO H_2_O. Quenching of endogenous peroxidase was performed by incubating slides for 20 minutes in the dark with 3% (v/v) hydrogen peroxidase in RO H_2_O, then rinsing slides in RO H_2_O twice for 5 minutes each and once in TBST (0.5M Tris, 9% w/v (w/v) NaCl, 0.5% (v/v) Tween-20, pH 7.6) for 5 minutes. Ki-67 (SP6) (Invitrogen, MA5-14520) was prepared in SignalStain® Antibody Diluent (CST, 8112) at 1:150 dilution and applied to slides for an overnight incubation in a humidified chamber at 4°C. The following day, slides were washed three times for 2 minutes each in TBST with agitation. Ki-67 was detected colorimetrically using the EnVision®+ System (DAB-based with HRP-conjugated antibodies [anti-Rabbit/Mouse mixture], DAKO, K4003), following the manufacturer’s protocol. Slides were counterstained in Mayer’s haematoxylin, blued, dehydrate, cleared, mounted and imaged as described above (see Histology).

### Quantitative image analysis

Histological colitis score (HCS) quantification (Figure 4.6B) was performed using the Aperio ImageScope v12.4.2.5010, following the method outlined by (Garcia-Hernandez et al., 2021). Briefly, three distinct layers were defined using different colours: total Swiss-roll length, areas of inflammation/injury and areas of erosion/ulceration. The “Pen” tool was used to manually outline the Swiss-roll, and the length of each category was measured. The total segment lengths for each region were summed, and the percentage of injury and ulceration relative to total length was calculated. HCS was determined using the following formula, based on the assumption that complete epithelial loss (i.e. erosion or ulceration) leads to maximal barrier dysfunction and worsened disease outcomes:

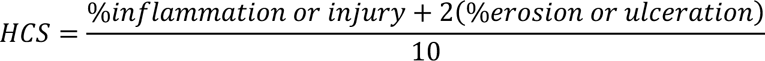

For goblet cell theca area quantification (Figure 5D), individual goblet cell theca areas were manually outlined using the “Pen” tool in Aperio ImageScope v12.4.2.5010. The average area per goblet cell was determined by measuring >100 goblet cells per animal.

For quantification of PAS-AB^+ve^ staining in the upper or lower crypts (Figure 5C), only fully visible crypts were included in the analysis. PAS-AB^+ve^ stained regions in the top or bottom half of the crypt were manually outlined and measured using the “Pen” tool in Aperio ImageScope.

For quantification of PAS-AB^+ve^ staining in the colon epithelium (Figure5B, E, F), analysis was performed using Fiji ImageJ. The areas of interest (i.e distal colon) were isolated by defining a Region of Interest (ROI). Colour deconvolution was applied using the “Colour Deconvolution 2” tool to separate PAS^+ve^, AB^+ve^, and haematoxylin staining, allowing mucin staining to be distinguished from background and nuclear staining. PAS^+ve^ and AB^+ve^ channels were then thresholded (Image>Adjust>Threshold (default)) to generate binary images. Watershed separation plugin (Process>Binary>Watershed) was applied to separate the structures. PAS^+ve^ or AB^+ve^ areas were quantified by analysing the structures of interest within the ROI (Analyze>Analyze Particle). The total PAS-AB^+ve^ mucin area was obtained by summing the quantified PAS^+ve^ and AB^+ve^ areas.

For quantification of immunofluorescence staining, Fiji ImageJ was used. Spatial calibration was first conducted by setting the appropriate scale for each image (Analyse>Set Scale). Quantification of the size of RAB5 vesicles (Figure 2C) was performed by manually drawing an ROI comprising of more than 5 contiguous intestinal epithelial cells displaying clear apical-basal orientation, while excluding non-specific or luminal debris staining in the organoid lumen, using the polygonal, freehand and line selection tools. The RAB5 channel was then thresholded to create a binary image containing the structures of interest (Image>Adjust>Threshold (default)) followed by the application of the watershed separation plugin (Process>Binary>Watershed) to separate the structures (referred to herein as particles). The particles within the ROI were then analysed (Analyze>Analyze Particle) to obtain the average size of particle.

Quantification of apical, lateral, basal and cytoplasmic fluorescence signals on whole-mount fluorescence images was conducted by manually drawing ROIs around the relevant structures using the polygonal, freehand and line selection tools and measured using the ‘Measure’ (Analyse>Measure) function. A minimum of 3 ROIs (comprising of >5 cells one after another) per stack, and at least 3 z-sections with clear apical-to-basal orientation, were analysed per organoid.

Co-localisation analysis was performed using the Just Another Colocalisation Plugin (JaCoP; Bioimaging and Optics Platform, BIOP). Otsu thresholding was applied to define object boundaries.

### Statistics and reproducibility

Numerical source data for all graphs are provided in Methods and Figure legends. Statistical tests were performed using GraphPad Prism 8 Software (GraphPad, San Diego, CA) *via* Student’s unpaired t-tests when comparing between two groups. One-way ANOVA or two-way ANOVA (with Tukey post-hoc comparisons) was used for multiple comparisons. Each mouse was assessed as an individual sample. All data were obtained by performing at least *n* > 2 independent experiments with representative data shown and expressed as the mean ± standard error of the mean (S.E.M). *P values* < 0.05 were considered as statistically significant. Significance levels were split further as follows: \*\**P* < 0.01, \*\*\**P* < 0.001, \*\*\*\**P* < 0.0001.

### Human research ethics and human intestinal biopsy collection

Human intestinal tissue collection for this study was approved by the Human Research Ethics Committee (Melbourne Health HREC/74911/MH-2021) of the Royal Melbourne Hospital, in accordance with the National Health and Medical Research Council (NHMRC) National Statement on Ethical Conduct in Human Research (2008) and the CPMP/ICH-135/95 Good Clinical Practice guidelines. Site-specific governance approvals were obtained from the Walter and Eliza Hall Institute (WEHI) and the University of Melbourne, with experimental analyses of all biopsies conducted at WEHI. Collaboration between all participating institutions was formalised through the Melbourne Academic Center for Health Research Collaboration Agreement (Non-Commercial). All research involving human participants was conducted in compliance with the principles outlined in the WMA Declaration of Helsinki and the Department of Health and Human Services Belmont Report. Written informed consent was obtained from all participants prior to sample collection.

Adult patients undergoing endoscopic evaluation (flexible sigmoidoscopy or colonoscopy) at the Royal Melbourne Hospital were screened for study inclusion ^74^. Patients were excluded if they had active systemic infections (gastrointestinal or non-gastrointestinal), malignancies undergoing active treatment, recent use of non-steroidal anti-inflammatory drugs (within one month), hereditary or familial polyposis syndromes, or non-IBD forms of colitis (including microscopic colitis, ischemic colitis, diversion colitis, or diverticulitis). Eligible patients were recruited by a gastroenterologist and provided signed informed consent. For patients diagnosed with inflammatory bowel disease (IBD), biopsies were collected endoscopically from inflamed, non-inflamed, and marginally inflamed intestinal regions. For non-IBD control patients, biopsies were obtained from non-inflamed intestinal segments following the same exclusion criteria. Intestinal biopsies were collected using Boston Scientific Radial Jaw™ biopsy forceps and Olympus EVIS EXERA III endoscopes, immediately placed in ice-cold DPBS (Thermo Fisher Scientific Cat#14190144) supplemented with protease inhibitors (Merck Cat#4693132001) and phosphatase inhibitors (Merck Cat#4906837001), and then homogenized with a stainless steel ball bearing in a Qiagen TissueLyzerII (30 Hz, 1 min) in 0.4 mL of ice-cold RIPA buffer (10 mM Tris— HCl pH 8.0, 1 mM EGTA, 2 mM MgCl_2_, 0.5% v/v Triton X-100, 0.1% w/v sodium deoxycholate, 0.5% w/v SDS, and 90 mM NaCl) supplemented with 1x Protease and Phosphatase Inhibitor Cocktail (Cell Signaling Technology Cat#5872) and 100 U/mL Benzonase (Sigma-Aldrich Cat#E1014). Lysates were stored at −80°C for downstream analysis *via* immunoblot.

## ACKNOWLEDGEMENTS

We acknowledge scholarship support for J.J. (La Trobe Graduate Research Scholarship and Full Fee Research Scholarship) and S.T. (La Trobe University Research Training Program Scholarship). We are grateful to the Australian Research Council for grant support to E.F.L., W.D.F., J.M.Mariadason (DP190102612); the National Health and Medical Research Council to A.L.S. (2002965), K.D. and A.S.Y. (2010704, 1136592), PDC (2008909), J.M.Murphy (1172929, 2034104, 9000719); the Victorian Government Operational Infrastructure Support Scheme; the Kenneth Rainin Foundation to J.M.Murphy, B.C., A.L.S. and A.H.A.-A; the US DOD (HT94252310088) to A.S.Y.; and the Victorian Cancer Agency to E.F.L. (MCRF19045) for fellowship support.

## AUTHOR CONTRIBUTIONS

J.J., S.T., designed, performed, and analysed experiments and wrote the paper; T.J.H., S.L.E., A.H.A-A, K.P., S.N.Y., M.E., D.B., C.M.R., R.N., L.J.J., designed, performed experiments, analysed and interpreted data; P.D.C., K.D., B.T.K., A.S.Y., J.M.M., B.C., A.L.S., J.M.M. analysed and interpreted data; W.D.F., E.F.L. designed the project, analysed, interpreted data, and wrote the paper. All authors commented on the manuscript.

## ADDITIONAL INFORMATION

Correspondence and requests for materials should be addressed to Erinna F. Lee or W. Douglas Fairlie.

## REFERENCES

1. Ellinghaus, D. et al. Genome-wide association analysis in primary sclerosing cholangitis and ulcerative colitis identifies risk loci at GPR35 and TCF4. Hepatology 58, 1074–1083 (2013).

2. Hampe, J. et al. A genome-wide association scan of nonsynonymous SNPs identifies a susceptibility variant for Crohn disease in ATG16L1. Nat Genet 39, 207–211 (2007).

3. Henckaerts, L. et al. Genetic variation in the autophagy gene ULK1 and risk of Crohn’s disease. Inflamm Bowel Dis 17, 1392–1397 (2011).

4. Lahiri, A., Hedl, M. & Abraham, C. MTMR3 risk allele enhances innate receptor-induced signaling and cytokines by decreasing autophagy and increasing caspase-1 activation. Proc Natl Acad Sci U S A 112, 10461–10466 (2015).

5. Lassen, K.G. et al. Genetic Coding Variant in GPR65 Alters Lysosomal pH and Links Lysosomal Dysfunction with Colitis Risk. Immunity 44, 1392–1405 (2016).

6. Morgan, A.R., Lam, W.J., Han, D.Y., Fraser, A.G. & Ferguson, L.R. Association Analysis of ULK1 with Crohn’s Disease in a New Zealand Population. Gastroenterol Res Pract 2012, 715309 (2012).

7. Parkes, M. et al. Sequence variants in the autophagy gene IRGM and multiple other replicating loci contribute to Crohn’s disease susceptibility. Nat Genet 39, 830–832 (2007).

8. Prescott, N.J. et al. A nonsynonymous SNP in ATG16L1 predisposes to ileal Crohn’s disease and is independent of CARD15 and IBD5. Gastroenterology 132, 1665–1671 (2007).

9. Rioux, J.D. et al. Genome-wide association study identifies new susceptibility loci for Crohn disease and implicates autophagy in disease pathogenesis. Nat Genet 39, 596–604 (2007).

10. Tran, S., Juliani, J., Fairlie, W.D. & Lee, E.F. The emerging roles of autophagy in intestinal epithelial cells and its links to inflammatory bowel disease. Biochem Soc Trans 51, 811–826 (2023).

11. Wellcome Trust Case Control, C. Genome-wide association study of 14,000 cases of seven common diseases and 3,000 shared controls. Nature 447, 661-678 (2007).

12. Benjamin, J.L., Sumpter, R., Jr., Levine, B. & Hooper, L.V. Intestinal epithelial autophagy is essential for host defense against invasive bacteria. Cell Host Microbe 13, 723–734 (2013).

13. Cabrera, S. et al. ATG4B/autophagin-1 regulates intestinal homeostasis and protects mice from experimental colitis. Autophagy 9, 1188–1200 (2013).

14. Cadwell, K. et al. A key role for autophagy and the autophagy gene Atg16l1 in mouse and human intestinal Paneth cells. Nature 456, 259–263 (2008).

15. Cadwell, K., Patel, K.K., Komatsu, M., Virgin, H.W.t. & Stappenbeck, T.S. A common role for Atg16L1, Atg5 and Atg7 in small intestinal Paneth cells and Crohn disease. Autophagy 5, 250–252 (2009).

16. Feng, Y., Wang, Y., Wang, P., Huang, Y. & Wang, F. Short-Chain Fatty Acids Manifest Stimulative and Protective Effects on Intestinal Barrier Function Through the Inhibition of NLRP3 Inflammasome and Autophagy. Cell Physiol Biochem 49, 190–205 (2018).

17. Lassen, K.G. et al. Atg16L1 T300A variant decreases selective autophagy resulting in altered cytokine signaling and decreased antibacterial defense. Proc Natl Acad Sci U S A 111, 7741–7746 (2014).

18. Naama, M. et al. Autophagy controls mucus secretion from intestinal goblet cells by alleviating ER stress. Cell Host Microbe 31, 433–446 e434 (2023).

19. Nighot, P.K. & Blikslager, A.T. Chloride channel ClC-2 modulates tight junction barrier function via intracellular trafficking of occludin. Am J Physiol Cell Physiol 302, C178–187 (2012).

20. Patel, K.K. et al. Autophagy proteins control goblet cell function by potentiating reactive oxygen species production. EMBO J 32, 3130–3144 (2013).

21. Saha, K. et al. Autophagy Reduces the Degradation and Promotes Membrane Localization of Occludin to Enhance the Intestinal Epithelial Tight Junction Barrier against Paracellular Macromolecule Flux. J Crohns Colitis 17, 433–449 (2023).

22. Wittkopf, N. et al. Lack of intestinal epithelial atg7 affects paneth cell granule formation but does not compromise immune homeostasis in the gut. Clin Dev Immunol 2012, 278059 (2012).

23. Zhao, Z. et al. Autophagosome-independent essential function for the autophagy protein Atg5 in cellular immunity to intracellular pathogens. Cell Host Microbe 4, 458–469 (2008).

24. Cooney, R. et al. NOD2 stimulation induces autophagy in dendritic cells influencing bacterial handling and antigen presentation. Nat Med 16, 90–97 (2010).

25. Martin, P.K. et al. Autophagy proteins suppress protective type I interferon signalling in response to the murine gut microbiota. Nat Microbiol 3, 1131–1141 (2018).

26. Saitoh, T. et al. Loss of the autophagy protein Atg16L1 enhances endotoxin-induced IL-1beta production. Nature 456, 264–268 (2008).

27. Sanjuan, M.A. et al. Toll-like receptor signalling in macrophages links the autophagy pathway to phagocytosis. Nature 450, 1253–1257 (2007).

28. Strisciuglio, C. et al. Impaired autophagy leads to abnormal dendritic cell-epithelial cell interactions. J Crohns Colitis 7, 534–541 (2013).

29. Zhang, H. et al. Myeloid ATG16L1 Facilitates Host-Bacteria Interactions in Maintaining Intestinal Homeostasis. J Immunol 198, 2133–2146 (2017).

30. Juliani, J. et al. BECLIN-1 is essential for the maintenance of gastrointestinal epithelial integrity by regulating endocytic trafficking, F-actin organization, and lysosomal function. Autophagy Reports 4, 2484494 (2025).

31. Tran, S. et al. BECLIN1 is essential for intestinal homeostasis involving autophagy-independent mechanisms through its function in endocytic trafficking. Commun Biol 7, 209 (2024).

32. Dong, L.W. et al. Prognostic significance of Beclin 1 in intrahepatic cholangiocellular carcinoma. Autophagy 7, 1222–1229 (2011).

33. Gong, C. et al. Beclin 1 and autophagy are required for the tumorigenicity of breast cancer stem-like/progenitor cells. Oncogene 32, 2261–2272, 2272e 2261-2211 (2013).

34. Liang, X.H. et al. Induction of autophagy and inhibition of tumorigenesis by beclin 1. Nature 402, 672–676 (1999).

35. Miracco, C. et al. Protein and mRNA expression of autophagy gene Beclin 1 in human brain tumours. Int J Oncol 30, 429–436 (2007).

36. Naguib, M. & Rashed, L.A. Serum level of the autophagy biomarker Beclin-1 in patients with diabetic kidney disease. Diabetes Res Clin Pract 143, 56–61 (2018).

37. Pickford, F. et al. The autophagy-related protein beclin 1 shows reduced expression in early Alzheimer disease and regulates amyloid beta accumulation in mice. J Clin Invest 118, 2190–2199 (2008).

38. Shibata, M. et al. Regulation of intracellular accumulation of mutant Huntingtin by Beclin 1. J Biol Chem 281, 14474–14485 (2006).

39. Valente, G. et al. Expression and clinical significance of the autophagy proteins BECLIN 1 and LC3 in ovarian cancer. Biomed Res Int 2014, 462658 (2014).

40. Bieri, G. et al. Proteolytic cleavage of Beclin 1 exacerbates neurodegeneration. Mol Neurodegener 13, 68 (2018).

41. Li, Z. et al. Genetic and epigenetic silencing of the beclin 1 gene in sporadic breast tumors. BMC Cancer 10, 98 (2010).

42. Qu, X., et al. Promotion of tumorigenesis by heterozygous disruption of the beclin 1 autophagy gene. J Clin Invest 112, 1809–1820 (2003).

43. Rohn, T.T. et al. Depletion of Beclin-1 due to proteolytic cleavage by caspases in the Alzheimer’s disease brain. Neurobiol Dis 43, 68–78 (2011).

44. Russo, R. et al. Calpain-mediated cleavage of Beclin-1 and autophagy deregulation following retinal ischemic injury in vivo. Cell Death Dis 2, e144 (2011).

45. Wang, J. et al. The long noncoding RNA H19 promotes tamoxifen resistance in breast cancer via autophagy. J Hematol Oncol 12, 81 (2019).

46. Zhou, L. et al. YTHDC1 inhibits autophagy-dependent NF-kappaB signaling by stabilizing Beclin1 mRNA in macrophages. J Inflamm (Lond) 21, 22 (2024).

47. Khan, S. et al. Cyclic GMP-AMP synthase contributes to epithelial homeostasis in intestinal inflammation via Beclin-1-mediated autophagy. FASEB J 36, e22282 (2022).

48. Ge, X. et al. The Loss of YTHDC1 in Gut Macrophages Exacerbates Inflammatory Bowel Disease. Adv Sci (Weinh) 10, e2205620 (2023).

49. DeRoche, T.C., Xiao, S.Y. & Liu, X. Histological evaluation in ulcerative colitis. Gastroenterol Rep (Oxf) 2, 178–192 (2014).

50. Matthew-Onabanjo, A.N. et al. Beclin 1 Promotes Endosome Recruitment of Hepatocyte Growth Factor Tyrosine Kinase Substrate to Suppress Tumor Proliferation. Cancer Res 80, 249–262 (2020).

51. McKnight, N.C. et al. Beclin 1 is required for neuron viability and regulates endosome pathways via the UVRAG-VPS34 complex. PLoS Genet 10, e1004626 (2014).

52. Noguchi, S. et al. Beclin 1 regulates recycling endosome and is required for skin development in mice. Commun Biol 2, 37 (2019).

53. Rohatgi, R.A. et al. Beclin 1 regulates growth factor receptor signaling in breast cancer. Oncogene 34, 5352–5362 (2015).

54. Wijshake, T. et al. Tumor-suppressor function of Beclin 1 in breast cancer cells requires E-cadherin. Proc Natl Acad Sci U S A 118 (2021).

55. Bryant, D.M. & Stow, J.L. The ins and outs of E-cadherin trafficking. Trends Cell Biol 14, 427–434 (2004).

56. Apodaca, G. Endocytic traffic in polarized epithelial cells: role of the actin and microtubule cytoskeleton. Traffic 2, 149–159 (2001).

57. Vasioukhin, V., Bauer, C., Yin, M. & Fuchs, E. Directed actin polymerization is the driving force for epithelial cell-cell adhesion. Cell 100, 209–219 (2000).

58. Woichansky, I., Beretta, C.A., Berns, N. & Riechmann, V. Three mechanisms control E-cadherin localization to the zonula adherens. Nat Commun 7, 10834 (2016).

59. Mizoguchi, A. Animal models of inflammatory bowel disease. Prog Mol Biol Transl Sci 105, 263–320 (2012).

60. Park, Y.H. et al. Adequate Dextran Sodium Sulfate-induced Colitis Model in Mice and Effective Outcome Measurement Method. J Cancer Prev 20, 260–267 (2015).

61. Reehorst, C.M. et al. EHF is essential for epidermal and colonic epithelial homeostasis, and suppresses Apc-initiated colonic tumorigenesis. Development 148 (2021).

62. Garcia-Hernandez, V. et al. Systematic Scoring Analysis for Intestinal Inflammation in a Murine Dextran Sodium Sulfate-Induced Colitis Model. J Vis Exp (2021).

63. Gustafsson, J.K. & Johansson, M.E.V. The role of goblet cells and mucus in intestinal homeostasis. Nat Rev Gastroenterol Hepatol 19, 785–803 (2022).

64. Buisine, M.P. et al. Mucin gene expression in intestinal epithelial cells in Crohn’s disease. Gut 49, 544–551 (2001).

65. Choi, W.T. et al. Hypermucinous, Goblet Cell-Deficient and Crypt Cell Dysplasias in Inflammatory Bowel Disease are Often Associated with Flat/Invisible Endoscopic Appearance and Advanced Neoplasia on Follow-Up. J Crohns Colitis 16, 98–108 (2022).

66. Gersemann, M. et al. Differences in goblet cell differentiation between Crohn’s disease and ulcerative colitis. Differentiation 77, 84–94 (2009).

67. Johansson, M.E. et al. Bacteria penetrate the normally impenetrable inner colon mucus layer in both murine colitis models and patients with ulcerative colitis. Gut 63, 281–291 (2014).

68. Kang, Y., Park, H., Choe, B.H. & Kang, B. The Role and Function of Mucins and Its Relationship to Inflammatory Bowel Disease. Front Med (Lausanne) 9, 848344 (2022).

69. Pullan, R.D. et al. Thickness of adherent mucus gel on colonic mucosa in humans and its relevance to colitis. Gut 35, 353–359 (1994).

70. Van der Sluis, M. et al. Muc2-deficient mice spontaneously develop colitis, indicating that MUC2 is critical for colonic protection. Gastroenterology 131, 117–129 (2006).

71. Xue, Y., Zhang, H., Sun, X. & Zhu, M.J. Metformin Improves Ileal Epithelial Barrier Function in Interleukin-10 Deficient Mice. PLoS One 11, e0168670 (2016).

72. Laroui, H. et al. Dextran sodium sulfate (DSS) induces colitis in mice by forming nano-lipocomplexes with medium-chain-length fatty acids in the colon. PLoS One 7, e32084 (2012).

73. Gregorieff, A. et al. The ets-domain transcription factor Spdef promotes maturation of goblet and paneth cells in the intestinal epithelium. Gastroenterology 137, 1333–1345 e1331-1333 (2009).

74. Chiou, S. et al. An immunohistochemical atlas of necroptotic pathway expression. EMBO Mol Med 16, 1717–1749 (2024).

75. Pang, J. et al. A necroptotic-to-apoptotic signaling axis underlies inflammatory bowel disease. bioRxiv, 2024.2011.2013.623307 (2024).

76. Yantiss, R.K. & Odze, R.D. Diagnostic difficulties in inflammatory bowel disease pathology. Histopathology 48, 116–132 (2006).

77. Hinnant, T., Ning, W. & Lechler, T. Compartment specific responses to contractility in the small intestinal epithelium. PLoS Genet 20, e1010899 (2024).

78. Allen, D.C., Hamilton, P.W., Watt, P.C. & Biggart, J.D. Architectural morphometry in ulcerative colitis with dysplasia. Histopathology 12, 611–621 (1988).

79. Richter, A., Yang, K., Richter, F., Lynch, H.T. & Lipkin, M. Morphological and morphometric measurements in colorectal mucosa of subjects at increased risk for colonic neoplasia. Cancer Lett 74, 65–68 (1993).

80. Rubio, C.A., Schmidt, P.T., Lang-Schwarz, C. & Vieth, M. Branching crypts in inflammatory bowel disease revisited. J Gastroenterol Hepatol 37, 440–445 (2022).

81. Tanaka, M. et al. Morphologic criteria applicable to biopsy specimens for effective distinction of inflammatory bowel disease from other forms of colitis and of Crohn’s disease from ulcerative colitis. Scand J Gastroenterol 34, 55–67 (1999).

82. Oliver, M.G. & Specian, R.D. Cytoskeleton of intestinal goblet cells: role of actin filaments in baseline secretion. Am J Physiol 259, G991–997 (1990).

83. Ehre, C. et al. Barrier role of actin filaments in regulated mucin secretion from airway goblet cells. Am J Physiol Cell Physiol 288, C46–56 (2005).

84. Mege, R.M. & Ishiyama, N. Integration of Cadherin Adhesion and Cytoskeleton at Adherens Junctions. Cold Spring Harb Perspect Biol **9** (2017).

85. Gaur, P. et al. Rab7-dependent regulation of goblet cell protein CLCA1 modulates gastrointestinal homeostasis. Elife 12 (2024).

86. Campbell, B.J., Yu, L.G. & Rhodes, J.M. Altered glycosylation in inflammatory bowel disease: a possible role in cancer development. Glycoconj J 18, 851–858 (2001).

87. Corfield, A.P. et al. Colonic mucins in ulcerative colitis: evidence for loss of sulfation. Glycoconj J 13, 809–822 (1996).

88. Larsson, J.M. et al. Altered O-glycosylation profile of MUC2 mucin occurs in active ulcerative colitis and is associated with increased inflammation. Inflamm Bowel Dis 17, 2299–2307 (2011).

89. McGuckin, M.A., Linden, S.K., Sutton, P. & Florin, T.H. Mucin dynamics and enteric pathogens. Nat Rev Microbiol 9, 265–278 (2011).

90. Raouf, A.H. et al. Sulphation of colonic and rectal mucin in inflammatory bowel disease: reduced sulphation of rectal mucus in ulcerative colitis. Clin Sci (Lond) 83, 623–626 (1992).

91. Van Klinken, B.J., Van der Wal, J.W., Einerhand, A.W., Buller, H.A. & Dekker, J. Sulphation and secretion of the predominant secretory human colonic mucin MUC2 in ulcerative colitis. Gut 44, 387–393 (1999).

92. Pullan, R.D. Colonic mucus, smoking and ulcerative colitis. Ann R Coll Surg Engl 78, 85–91 (1996).

93. Strugala, V., Dettmar, P.W. & Pearson, J.P. Thickness and continuity of the adherent colonic mucus barrier in active and quiescent ulcerative colitis and Crohn’s disease. Int J Clin Pract 62, 762–769 (2008).

94. Fasseu, M. et al. Identification of restricted subsets of mature microRNA abnormally expressed in inactive colonic mucosa of patients with inflammatory bowel disease. PLoS One 5 (2010).

95. Hao, X., Yang, B., Liu, X., Yang, H. & Liu, X. Expression of Beclin1 in the colonic mucosa tissues of patients with ulcerative colitis. Int J Clin Exp Med 8, 21098–21105 (2015).

96. Wang, S.L. et al. Intestinal autophagy links psychosocial stress with gut microbiota to promote inflammatory bowel disease. Cell Death Dis 10, 391 (2019).

97. Zhu, H. et al. Regulation of autophagy by a beclin 1-targeted microRNA, miR-30a, in cancer cells. Autophagy 5, 816–823 (2009).

98. Levine, B., Mizushima, N. & Virgin, H.W. Autophagy in immunity and inflammation. Nature 469, 323–335 (2011).

99. Bray, K. et al. Autophagy suppresses RIP kinase-dependent necrosis enabling survival to mTOR inhibition. PLoS One 7, e41831 (2012).

100. Fairlie, W.D., Tran, S. & Lee, E.F. Crosstalk between apoptosis and autophagy signaling pathways. Int Rev Cell Mol Biol 352, 115–158 (2020).

101. Hou, W., Han, J., Lu, C., Goldstein, L.A. & Rabinowich, H. Autophagic degradation of active caspase-8: a crosstalk mechanism between autophagy and apoptosis. Autophagy 6, 891–900 (2010).

102. Lucin, K.M. et al. Microglial beclin 1 regulates retromer trafficking and phagocytosis and is impaired in Alzheimer’s disease. Neuron 79, 873–886 (2013).

103. Seo, J. et al. Beclin 1 functions as a negative modulator of MLKL oligomerisation by integrating into the necrosome complex. Cell Death Differ 27, 3065–3081 (2020).

104. Thorburn, J. et al. Autophagy controls the kinetics and extent of mitochondrial apoptosis by regulating PUMA levels. Cell Rep 7, 45–52 (2014).

105. Ferguson, L.R. et al. Genetic factors in chronic inflammation: single nucleotide polymorphisms in the STAT-JAK pathway, susceptibility to DNA damage and Crohn’s disease in a New Zealand population. Mutat Res 690, 108–115 (2010).

106. Miao, L.J. et al. Stat3 inhibits Beclin 1 expression through recruitment of HDAC3 in nonsmall cell lung cancer cells. Tumour Biol 35, 7097–7103 (2014).

107. Tang, Y., Tan, S.A., Iqbal, A., Li, J. & Glover, S.C. STAT3 Genotypic Variant rs744166 and Increased Tyrosine Phosphorylation of STAT3 in IL-23 Responsive Innate Lymphoid Cells during Pathogenesis of Crohn’s Disease. J Immunol Res 2019, 9406146 (2019).

108. Cho, D.H. et al. Caspase-mediated cleavage of ATG6/Beclin-1 links apoptosis to autophagy in HeLa cells. Cancer Lett 274, 95–100 (2009).

109. Wirawan, E. et al. Caspase-mediated cleavage of Beclin-1 inactivates Beclin-1-induced autophagy and enhances apoptosis by promoting the release of proapoptotic factors from mitochondria. Cell Death Dis 1, e18 (2010).

110. Patankar, J.V. & Becker, C. Cell death in the gut epithelium and implications for chronic inflammation. Nat Rev Gastroenterol Hepatol 17, 543–556 (2020).

111. el Marjou, F., et al. Tissue-specific and inducible Cre-mediated recombination in the gut epithelium. Genesis 39, 186–193 (2004).

112. Dekkers, J.F. et al. High-resolution 3D imaging of fixed and cleared organoids. Nat Protoc 14, 1756–1771 (2019).

