## Supplemental Material for "Sufficient levels of BECLIN1 are required for intestinal epithelial cell homeostasis and protection against unwanted intestinal inflammation"

Supplementary Figure 1

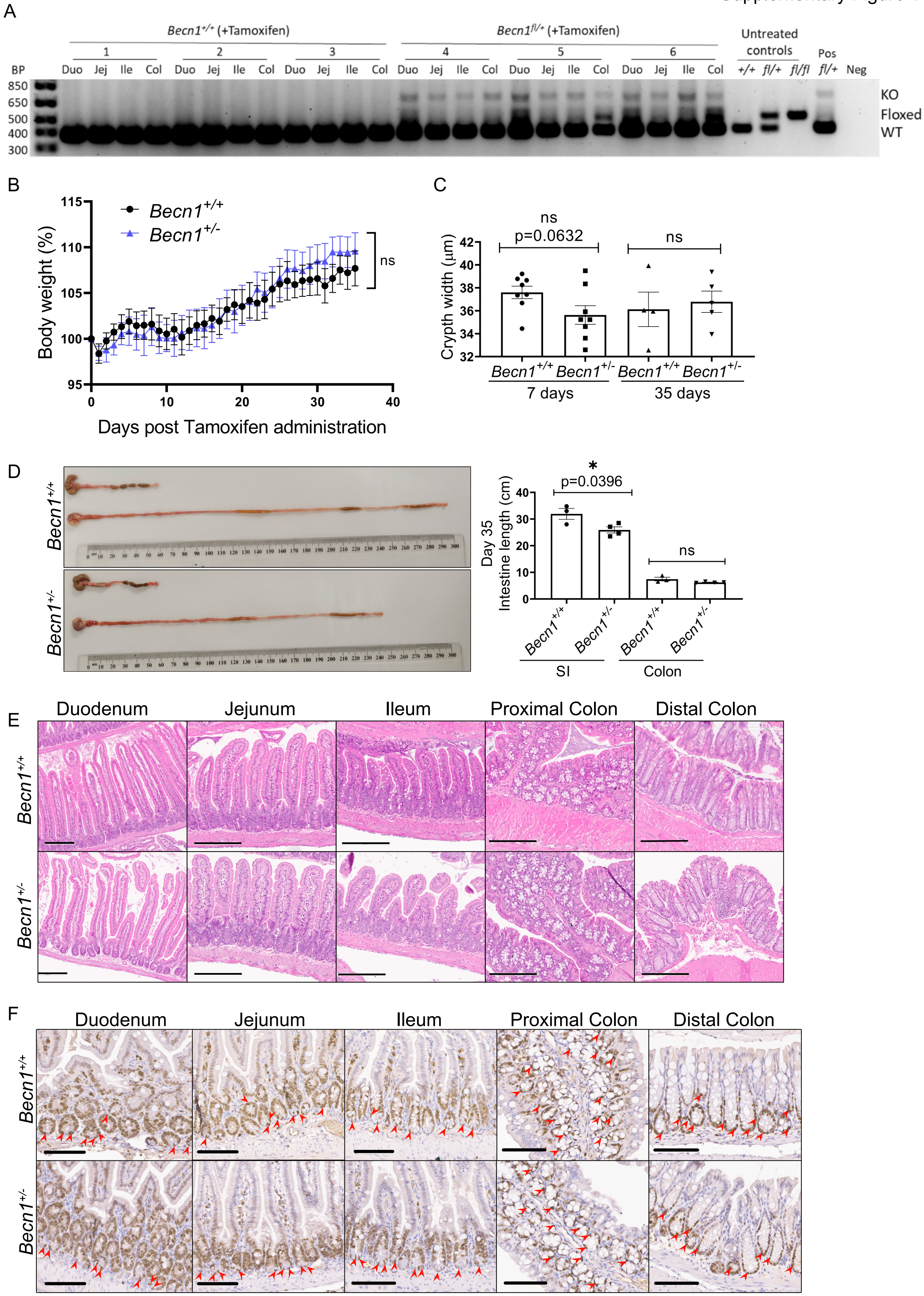

Supplementary Figure 2

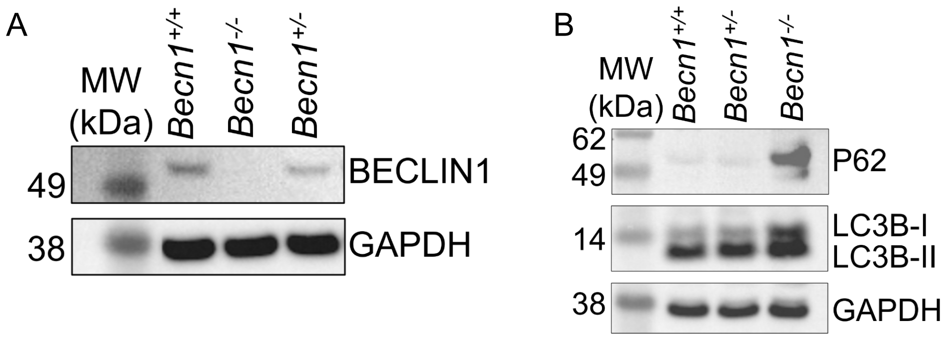

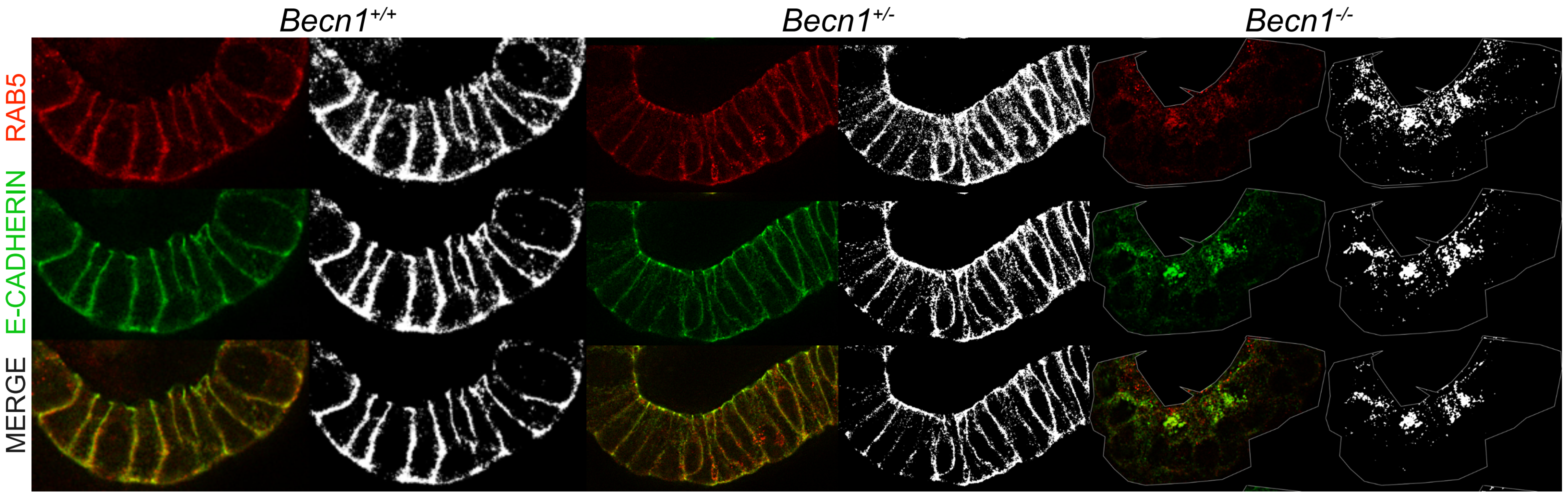
*Becn1*+/+
*Becn1*+/-
*Becn1*-/-
RAB5
E-CADHERIN
MERGE

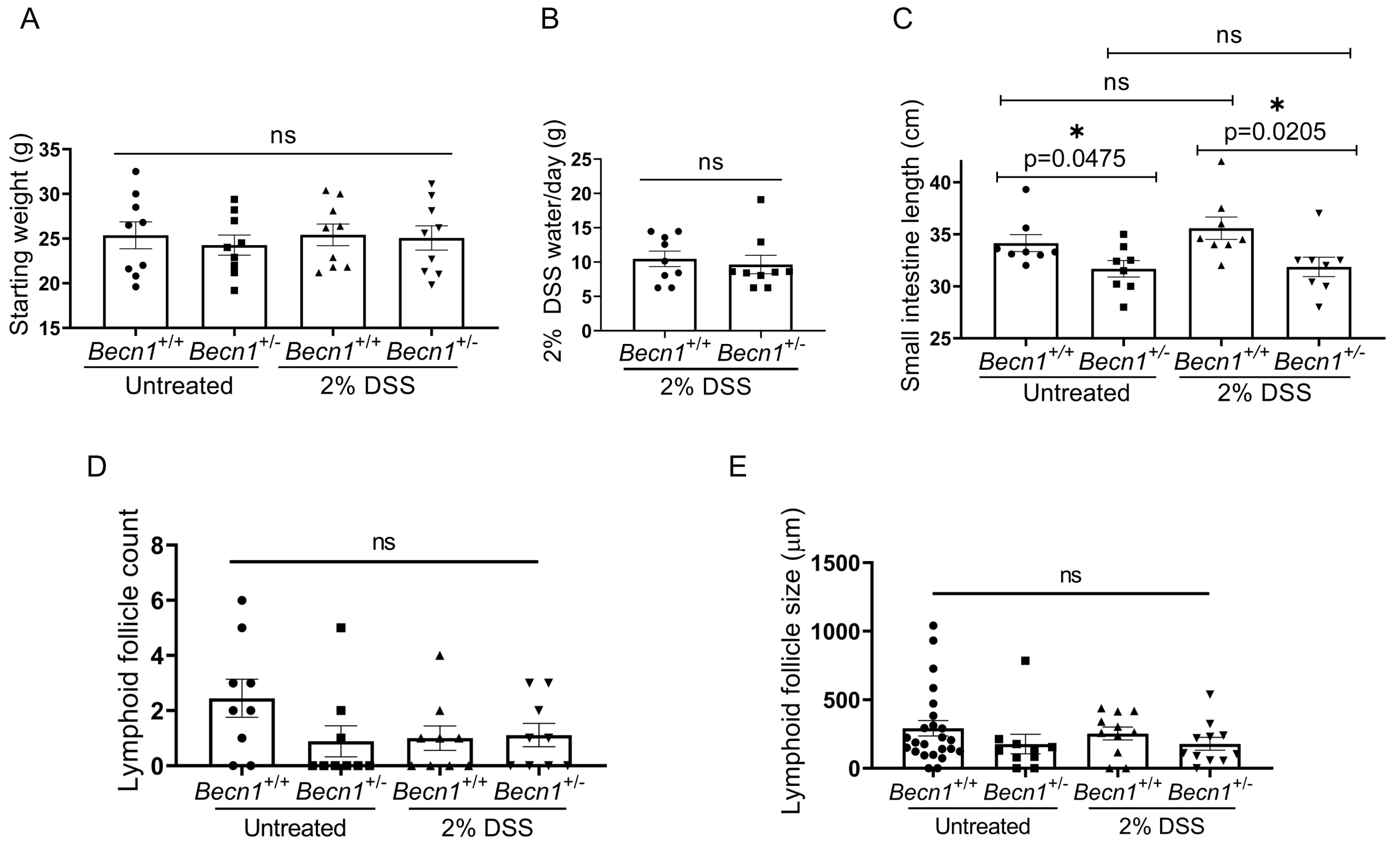
A
Starting weight (g)
ns
Becn1+/+ Becn1+/- Becn1+/+ Becn1+/-
Untreated
2% DSS
B
2% DSS water/day (g)
ns
Becn1+/+ Becn1+/-
2% DSS
C
Small intestine length (cm)
ns
ns
*
p=0.0475
*
p=0.0205
Becn1+/+ Becn1+/- Becn1+/+ Becn1+/-
Untreated
2% DSS
D
Lymphoid follicle count
ns
Becn1+/+ Becn1+/- Becn1+/+ Becn1+/-
Untreated
2% DSS
E
Lymphoid follicle size (μm)
ns
Becn1+/+ Becn1+/- Becn1+/+ Becn1+/-
Untreated
2% DSS

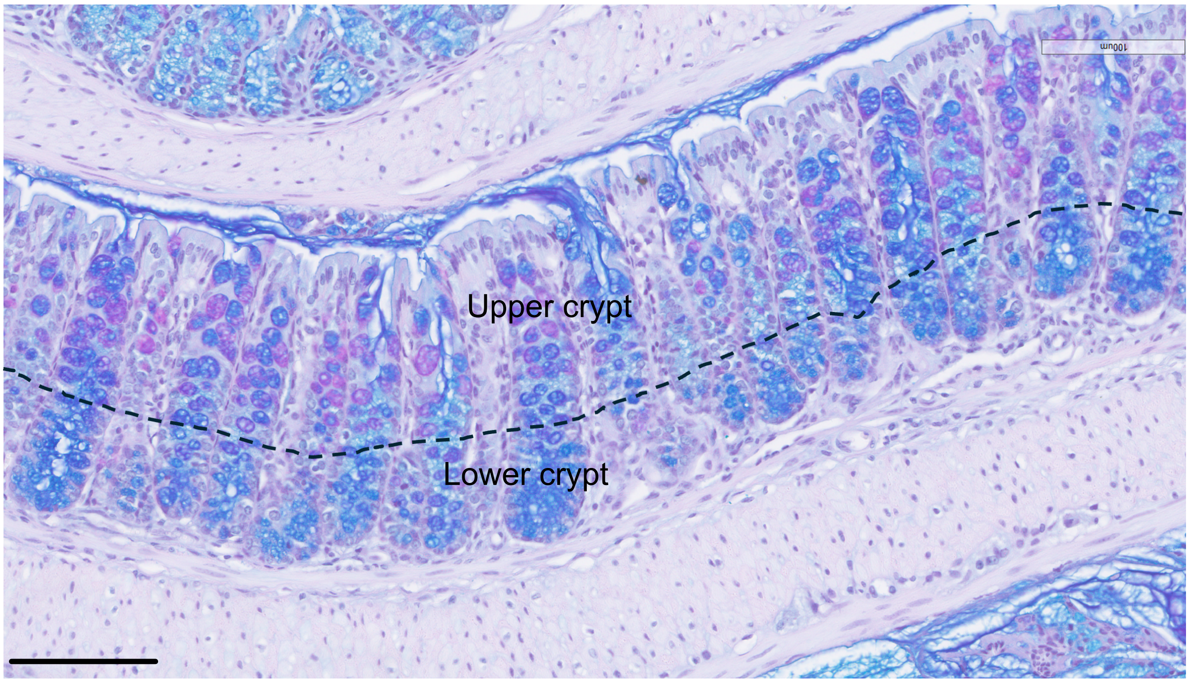
Upper crypt
Lower crypt

**Supplementary Figure 1. Monoallelic *Becn1* deletion in mice results in shortened small intestines and reduced colonic crypt length even when aged up to one month. (A)** Representative PCR genotyping of IECs isolated from different sections of the gastrointestinal tract of both *Becn1*$^{+/+}$*;Vil1-CreERT2*$^{Cre/+}$ and *Becn1*$^{fl/+}$*;Vil1-CreERT2*$^{Cre/+}$ mice two weeks post-Tamoxifen treatment. Numbers (1-6) correspond to individual animals. Untreated IEC controls (*+/+, fl/+* and *fl/fl*) were isolated from the duodenum of *Becn1*$^{+/+}$*; Becn1*$^{fl/+}$*;* and *Becn1*$^{fl/fl}$ *;Vil1-CreERT2*$^{Cre/+}$ mice without Tamoxifen treatment. Heterozygous *Becn1* deletion (Pos *fl/+*) is evidenced by the presence of a 398 bp band corresponding to the 'wild-type (WT)' allele, the loss or significant reduction of the 510 bp 'Floxed' band, and the emergence of a 721 bp band corresponding to the deleted allele (KO). **(B)** Body weight of mice over time, normalised to day 0. Data represent *n* > 6 biologically independent mice of each genotype from *n* = 3 independent experiments. **(C)** Distal colon crypt width of *Becn1*$^{+/+}$ and *Becn1*$^{+/-}$ mice at days 7 and 35 post-tamoxifen administration. Data represents *n* > 4 animals per genotype from *n* = 3 independent experiments. **(D)** Representative images of intestinal tracts of *Becn1*$^{+/+}$ and *Becn1*$^{+/-}$ mice along with the measurements of intestinal length at Day 35 post Tamoxifen administration. Data represents *n* > 3 animals per genotype. **(E)** H&E stained FFPE sections of *Becn1*$^{+/+}$ and *Becn1*$^{+/-}$ mice intestinal tract at 35 days post-tamoxifen administration. Scale bars represent 200 µm. **(F)** Representative KI-67 immunostaining of FFPE sections from *Becn1*$^{+/+}$ and *Becn1*$^{+/-}$ mice at day 7 post-tamoxifen administration across different segments of the small and large intestine. Red arrowheads indicate intact proliferative activity within the intestinal stem cell compartments. Scale bar = 100 µm. Data represents *n* = 6 from *n* = 3 independent experiments. Graphs show the mean ± S.E.M. Statistical significance was determined using unpaired (Student's) t-test. SI: small intestine. Duo: Duodenum. Jej: jejunum. Ile: ileum. Col: colon. BP: base pairs. Pos: positive control. Neg: negative control.

**Supplementary Figure 2. BECLIN1 reduction in intestinal organoids does not inhibit basal autophagic function. (A)** Representative Western immunoblot measuring BECLIN1 levels in *Becn1*$^{+/+}$*, Becn1*$^{+/-}$ *and Becn1*$^{-/-}$ organoids at day 7 post-4HT treatment. **(B)** Western blot assessment of autophagy flux in *Becn1*$^{+/+}$*, Becn1*$^{+/-}$ and *Becn1*$^{-/-}$ organoids, analysing total levels of P62, LC3B-1 and LC3B-II. Images are representative from *n* = 3 independent experiments with *n* = 3 different biological replicates.

**Supplementary Figure 3. Visualisation of RAB5 and E-CADHERIN colocalisation in intestinal organoids.** Representative images of whole-mount immunostained intestinal organoids from *Becn1*$^{+/+}$, *Becn1*$^{+/-}$ and *Becn1*$^{-/-}$ organoids showing RAB5 and E-CADHERIN staining, analysed using BioP-JACoP colocalisation with Otsu thresholding. Colocalised regions between RAB5 and E-CADHERIN are visualised in the MERGE (white) channel.

**Supplementary Figure 4. Baseline characteristics and intestinal length measurements of *Becn1*$^{+/+}$ and *Becn1*$^{+/-}$ mice following DSS or control treatment (A)** Starting weights of mice used in this experiment, measured at day zero prior to Tamoxifen injections. **(B)** The average amount of daily 2% DSS water intake, obtained by calculating the difference in weights of drinking water bottles at the start and end of experiment and dividing by the number of days the animals received treatment. **(C)** Small intestinal lengths of *Becn1*$^{+/+}$ and *Becn1*$^{+/-}$ mice who received normal drinking water (untreated) or 2% DSS drinking water (2% DSS) at endpoint. Data are representative of at least $n$ = 9 biologically independent mice from $n$ = 3 independent experiments. Graphs indicate the ± S.E.M. Statistical significance was determined using ordinary one-way ANOVA except in (B) where unpaired (Student's) t-test was used. **(D)** Number and **(E)** size of lymphoid follicles in the colon of *Becn1*$^{+/+}$ and *Becn1*$^{+/-}$ mice receiving normal drinking water or 2% DSS drinking water, calculated using H&E stained FFPE colon sections. Data are representative of at least $n$ = 9 biologically independent mice from $n$ = 3 independent experiments. Graphs indicate the ± S.E.M. Statistical significance was determined using ordinary one-way ANOVA except in (B) where unpaired (Student's) t-test was used.

**Supplementary Figure 5.** Representative PAS-AB-stained distal colonic epithelium, segmented into upper and lower crypt regions as indicated by dashed line for mucin quantification. Scale bar = 100 μm.
